# Notch signaling in germ line stem cells controls reproductive aging in *C. elegans*

**DOI:** 10.1101/2022.03.04.482923

**Authors:** Zuzana Kocsisova, Elena D. Bagatelas, Jesus Santiago-Borges, Hanyue Cecilia Lei, Brian M. Egan, Matthew C. Mosley, Daniel L. Schneider, Tim Schedl, Kerry Kornfeld

## Abstract

Reproductive aging in females often occurs early in life, resulting in a substantial post-reproductive lifespan. Despite the medical importance of age-related infertility, relatively little is known about mechanisms that control this age-related decline. *C. elegans* is a leading system for aging biology due to its short lifespan and powerful experimental tools, and detailed descriptions of molecular and cellular changes in the gonad during reproductive aging were recently reported. Here we show that reproductive aging occurs early in life in multiple species in the genus *Caenorhabditis*, indicating this is a feature of both female/male and hermaphrodite/male species. In mutants previously established to display delayed reproductive aging (*daf-2, eat-2*, phm-2), we observed correlations between changes in the distal germline and changes in egg-laying, consistent with the model that distal germline changes are a cause of reproductive aging. By screening for additional mutants that delay reproductive aging, we identified an allele of *che-3* with impaired sensory perception that displayed increased progeny production in mid-life, a pattern of reproductive aging distinct from previous mutants. To directly test the role of Notch signaling in the distal germline, we analyzed the effect of ectopic expression of the Notch effector gene SYGL-1. Ectopic expression of SYGL-1 was sufficient to delay reproductive aging, suggesting that an age-related decline in Notch signaling in the distal germline is a root cause of reproductive aging.

## Introduction

As adult animals advance through time, they display progressive degenerative changes in multiple organ systems. Most aging research focuses on the decline of life support systems that result in death and determine adult lifespan. However, there is growing appreciation that the age-related decline of the reproductive system, which results in infertility, is an important dimension of aging (Scharf et al., 2021). Because female reproduction typically declines well before the maximum adult lifespan, reproductive aging offers the possibility to experimentally investigate an aging organ in the context of a relatively intact animal. By contrast, studies of somatic aging are confounded by the simultaneous degeneration of multiple life support systems and the resulting cascade of failure. In addition to practical considerations, reproductive aging is conceptually important because progeny production is the ultimate goal of animal life. Thus, reproductive aging is a central issue in evolutionary theories of aging. Furthermore, reproductive aging is clinically relevant in humans. Age-related infertility is increasingly a concern for women who wait until middle-age to start families, and infertility treatments are a major source of medical expenses (Crawford and Steiner, 2015).

The nematode worm *Caenorhabditis elegans* is an experimentally powerful model for reproductive aging that is relevant to mammals (Athar and Templeman, 2022; Scharf et al., 2021). From a fertilized egg, the *C. elegans* hermaphrodite develops its entire soma and both arms of its reproductive tract, then begins laying eggs at ∼65 hours at 20°C (Byerly et al., 1976). At the peak of reproduction, oocytes are ovulated every ∼23 min (McCarter et al., 1999), resulting in an average of ∼150 progeny per day. Even when sperm are not limiting, reproduction ceases relatively early in life; in wild-type hermaphrodites with sufficient sperm from mating to males, the reproductive span is ∼9 days and the lifespan is ∼16 days (Pickett et al., 2013). Further, progeny production declines from a peak of ∼150 progeny per day on adult day 2 to ∼40 progeny per day on adult day 5 and to ∼12 progeny per day on adult day 7, while the animals are all still alive, moving, and feeding (Kocsisova et al., 2019). Because *C. elegans* has a self-fertile hermaphrodite/male sexual system (protandrous, androdioecious), it is an open question whether it has evolved an unusual type of reproductive aging compared to the more prevalent female/male (gonochoristic) species.

Because *C. elegans* is a premier system for genetic analysis, many mutations have been identified that delay age-related degenerative changes of somatic tissues and extend mean lifespan. Interestingly, only a small subset of these mutations have been reported to delay reproductive aging (Andux and Ellis, 2008; Angeles-Albores et al., 2017; Gems et al., 1998; Hughes et al., 2007, 2011; Luo et al., 2009; Qin and Hubbard, 2015; Templeman et al., 2018, 2020; Wang et al., 2014). A mutation of daf-2 that impairs the activity of the insulin/IGF-1 signaling pathway reduces the level of early progeny production, extends the reproductive span, and increases the level of progeny production late in life. Similar patterns are caused by *eat-2* and *phm-2* mutations that result in pathogenic infections and food avoidance behavior that drives dietary restriction (Kumar et al., 2019). These observations raise the question of whether reduced levels of early progeny production are the cause of delayed reproductive aging or an independent, correlated effect. Because the identification of mutations that delay reproductive aging is an important first step in elucidating the mechanisms of this process, the discovery of additional mutations that cause this phenotype is an important objective.

A detailed description of the molecular and cellular changes that occur during reproductive aging is important for generating hypotheses about the sequence of events that cause these changes (Scharf et al., 2021). This description is an essential foundation to comprehensively understand how genetic mutations influence this phenotype. Kocsisova et al. (2019) reasoned that because the most striking decline in reproduction (in hermaphrodites with sufficient sperm) occurs at approximately day 5 of adulthood, when mated hermaphrodites produce only ∼40 progeny - a mere quarter of peak reproduction - it would be relevant to analyze age-related changes in the germline that occur a day or two before this time. The decline in progeny production is preceded by decreased Notch signaling in the distal germ line, a decreased stem cell pool, a slower stem cell cycle, and decreased meiotic entry from the progenitor zone. Based on these observational studies, Kocsisova et al. (2019) hypothesized that these age-related molecular and cellular changes cause reproductive aging. However, this hypothesis has not been rigorously evaluated.

Here we show that reproductive aging occurs relatively early in two female/male species in the genus *Caenorhabditis*, suggesting this is a general feature and not specific to self-fertile hermaphrodites. To test the hypothesis that age-related changes in the distal germline are a cause of reproductive aging, we analyzed *daf-2, eat-2*, and *phm-2* mutants. Delayed changes in the distal germline were well correlated with extended progeny production, supporting the hypothesis that these changes are causal. To identify additional mutations that influence reproductive aging, we conducted a candidate screen and identified a *che-3* mutation that causes a pattern of delay in reproductive aging that is distinct from previously identified mutations. These *che-3* mutants have normal levels of early reproduction and an increase of midlife progeny production. This mutant illustrates that reduced early reproduction is not essential for delayed reproductive aging. Finally, we directly manipulated the Notch signaling pathway by increasing the expression levels of *sygl-1*, a direct Notch target gene that specifies the stem cell fate. Ectopic expression of *sygl-1* was sufficient to increase midlife reproduction without reducing the level of early reproduction or extending lifespan. These findings directly link Notch signaling output through SYGL-1 to reproductive aging, and we hypothesize that age-related changes in Notch signaling levels are an underlying cause of reproductive aging.

## Results

### Rapid reproductive aging occurs in hermaphrodite/male and female/male species in the genus *Caenorhabditis*

Hughes et al. (2007) showed that reproductive aging occurs rapidly compared to somatic aging in the species *C. elegans* that is hermaphrodite/male. To determine if rapid reproductive aging is specific to hermaphrodite/male nematode species, we analyzed *Caenorhabditis* species that are female/male. *Caenorhabditis* is well suited for comparative studies, since the genus contains approximately 60 described species (Dayi et al., 2021; Stevens et al., 2019). At least three of these are hermaphrodite/male (*C. elegans, C. briggsae*, and *C. tropicalis*), while the remaining species exhibit the ancestral female/male reproductive strategy (e.g. *C. brenneri, C. remanei*, and *C. angaria*) (Kiontke et al., 2011). We measured lifespan and progeny production of females of wild-type *C. brenneri* and *C. remanei*, as well as wild-type and *fog-2(lf)* mutant *C. elegans*, where the germline is feminized in hermaphrodites but males are unaffected. Females were mated to conspecific males. All three species displayed a peak of reproduction at days 2-3 of adulthood, followed by a decline in reproduction (**Figure 1**). Very few progeny were produced after day 8 (**Table S1**). By contrast, survival of animals was above 75% until day 10 (**Figure 1**). Thus, mated females from both female/male and hermaphrodite/male species in the genus *Caenorhabditis* displayed age-related reproductive decline several days before animals began to display age-related death.

**Figure 1:**
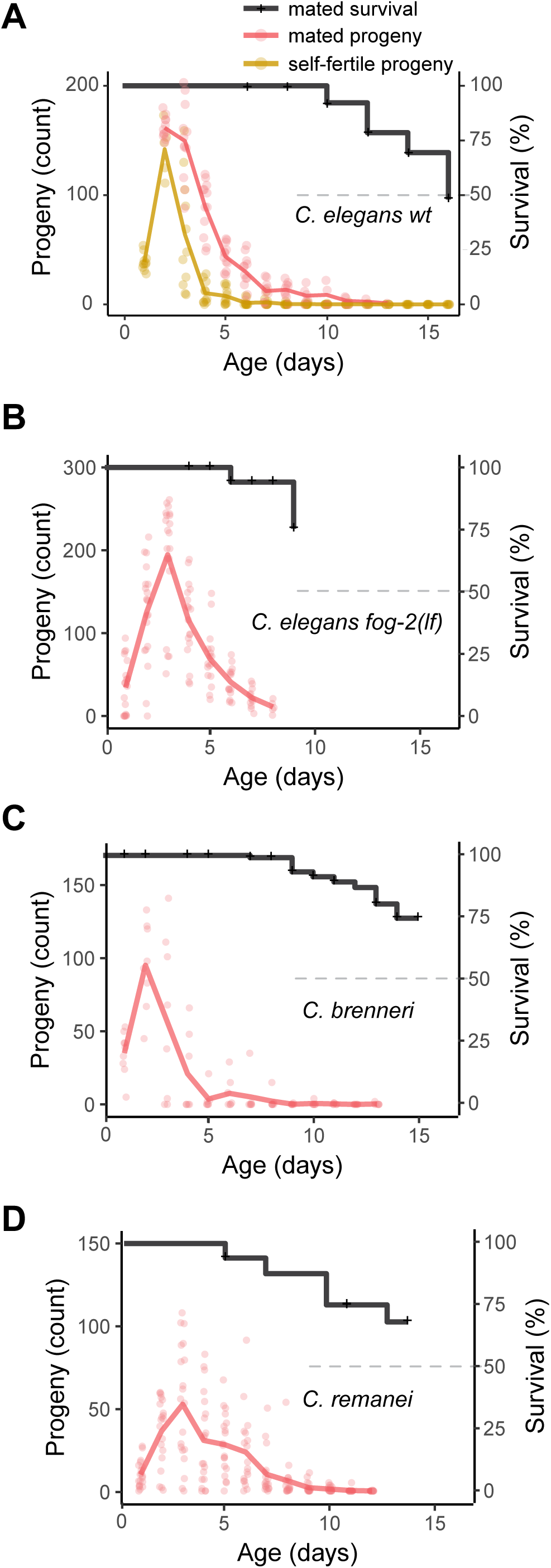
*C. elegans, C. brenneri* and *C. remanei* displayed rapid reproductive aging. The left axis indicates daily progeny production of unmated, self-fertile hermaphrodites (yellow), mated hermaphrodites (red), or mated females (red). Circles are values from one individual, and lines indicate the average. The right axis indicates survival of mated hermaphrodites or mated females (black). Vertical marks on survival plots indicate timepoints when animal(s) were censored due to matricidal hatching or crawling off the dish. In some experiments, the survival curve did not reach zero due to censoring. Dashed line indicates 50% survival. The horizontal axis indicates adult age in days. A) wild-type *C. elegans* strain N2 B) *C. elegans* fog-2(oz40) strain BS553. C) *C. brenneri* strain VX02253. D) *C. remanei* strain SB146. Panel A reproduced with permission from Kocsisova et al., (2019).

### Partial loss of insulin/IGF-1 signaling caused by *daf-2(e1370)* delayed reproductive aging and age-related changes in germ cells

To investigate the mechanism of rapid reproductive aging, we analyzed mutant strains that display delayed reproductive aging in hermaphrodites with sufficient sperm. *daf-2(e1370)* is a partial loss-of-function mutation in the insulin/IGF-1 receptor that delays somatic and reproductive aging (Hughes et al., 2007; Kenyon et al., 1993). We counted daily progeny production and analyzed three phases: early (day 1-4), mid-life (day 5-7), and late (day 8-15). Early progeny production by mated *daf-2(e1370)* hermaphrodites (180±30) was significantly lower than wild-type hermaphrodites (400±90), whereas mid-life progeny production was slightly but significantly higher than wild type (130±25 versus 100±30), and late progeny production was significantly higher than wild type (60±30 versus 10±10) (**Figure 2A-C, S1A, Table S2**). The total brood size of *daf-2(e1370)* hermaphrodites (360±70) was significantly lower than wild-type hermaphrodites (490±120) (**Figure 2B, Table S2**). Thus, daf-2 is necessary to promote high levels of early progeny production and rapid reproductive aging.

**Figure 2:**
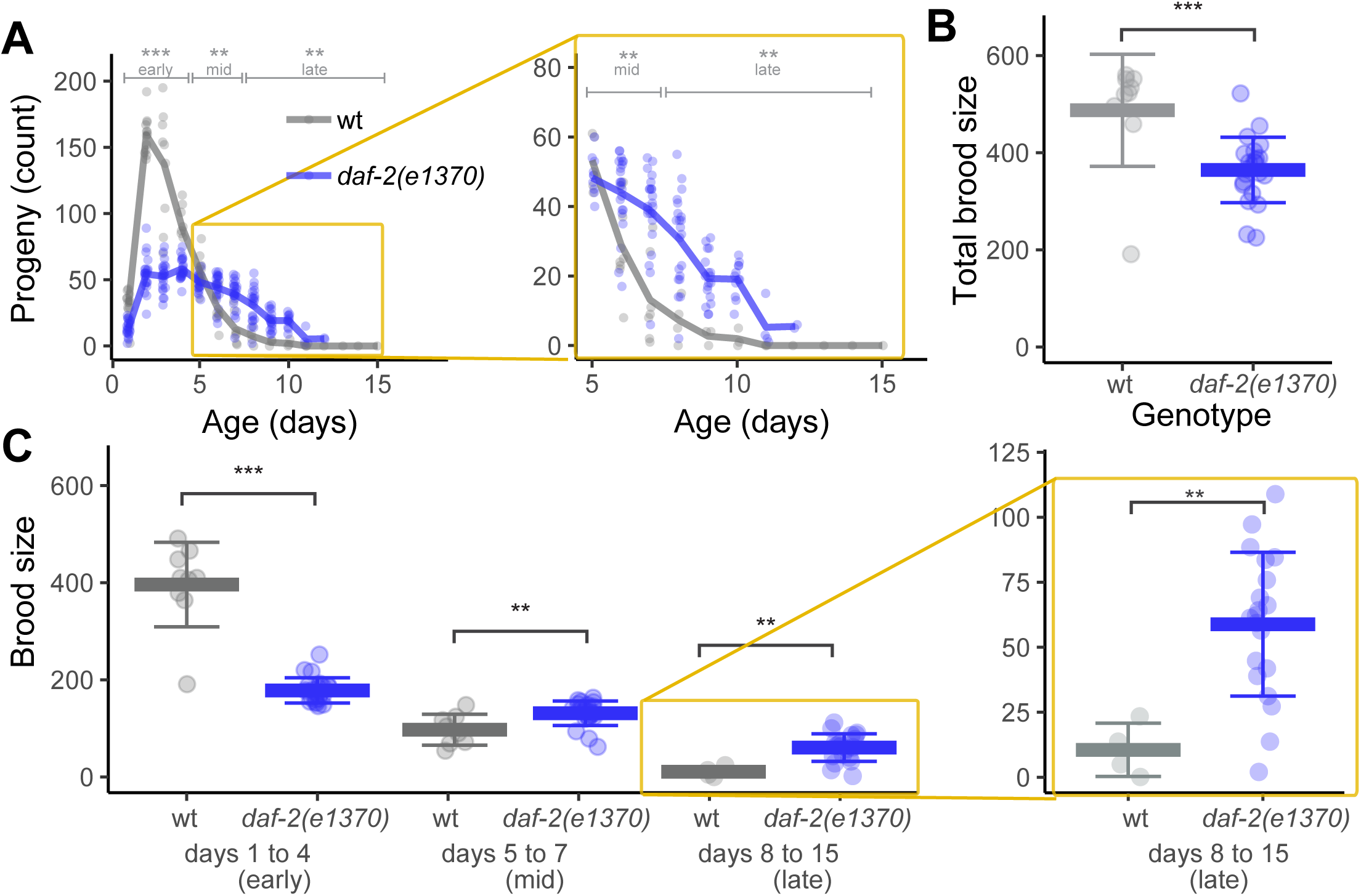
The *daf-2(e1370)* mutation delayed reproductive aging. Wild type (gray) and *daf-2(e1370)* (blue) hermaphrodites were mated to males. A) Lines are average daily progeny production, and points are data for individuals. Brackets above indicate early (days 1-4), mid-life (days 5-7) and late (days 8-15) progeny production periods. Within each time period, wild-type and mutant progeny totals were compared by a Wilcoxon Rank Sum test. The area in the yellow box is enlarged on the right. B) The total brood size, sum of progeny on days 1-15, compared by a Wilcoxon Rank Sum test. Bar and whiskers indicate mean ± SD. C) The early, mid-life, and late brood size. Kruskal–Wallis test. The area in the yellow box is enlarged on the right. * P<0.05, ** P<.001, *** P<.0001.

To determine if delayed reproductive aging is a general feature of lifespan extending mutations in the insulin/IGF-1 pathway, we analyzed additional alleles. Late progeny production was not significantly different in mated hermaphrodites of daf-2(e1368) (28±13), age-1(hx546) (11±7), or age-1(am88) (23±18) compared to wild type (22±15) (**Figure S1B-C, Table S2**). The total mated brood size of daf-2(e1368) (420±70) was significantly smaller than wild type (485±120) **(Figure S1D-F, Table S2**). Hughes (2005) reported that mated hermaphrodites of daf-2(m41) and age-1(hx546) did not display increased progeny production after day 9 (1.3±0.7 and 0.5±0.2, respectively) compared to wild-type (1.8±0.5). Thus, even among insulin/IGF-1 pathway mutations that extend longevity, the *daf-2(e1370)* allele is unusual in causing a delay of reproductive aging.

To determine the molecular and cellular changes in *daf-2(e1370)* hermaphrodites that may be responsible for delayed reproductive aging, we used the techniques described in Kocsisova et al., (2019) to measure germline phenotypes in day 1, 3, 5, and 7 mated hermaphrodites. To measure the distal-proximal extent of Notch signaling, we used an anti-FLAG antibody to stain *sygl-1(am307flag)* hermaphrodites, which express from the native locus SYGL-1 fused to 3xFLAG epitope. *sygl-1* is a direct target of Notch (Chen et al., 2020; Kershner et al., 2014; Shin et al., 2017) and thus a readout of the extent of Notch signaling output in the distal germline. We counted the number of cell diameters (cd, a measure of length) from the distal tip of the gonad to the last row with half or more FLAG immunoreactive cells. daf-2(+);*sygl-1(am307flag)* hermaphrodite controls displayed an age-related decrease in the extent of the SYGL-1 from 10±2 cd at day 1, to 7.5±2 cd at day 3, to 7±2 cd at day 5 (Kocsisova et al., 2019). daf-2(e1370);*sygl-1(am307flag)* hermaphrodites displayed a significantly longer SYGL-1 zone at day 1 (12±1.5 cd) and day 3 (9±1 cd) compared to these controls. Day 5 displayed a higher value (8.4±3 cd) that did not reach statistical significance with this sample size (P=0.15) (**Figure 3A-D, S1I-J, Table S3**).

**Figure 3:**
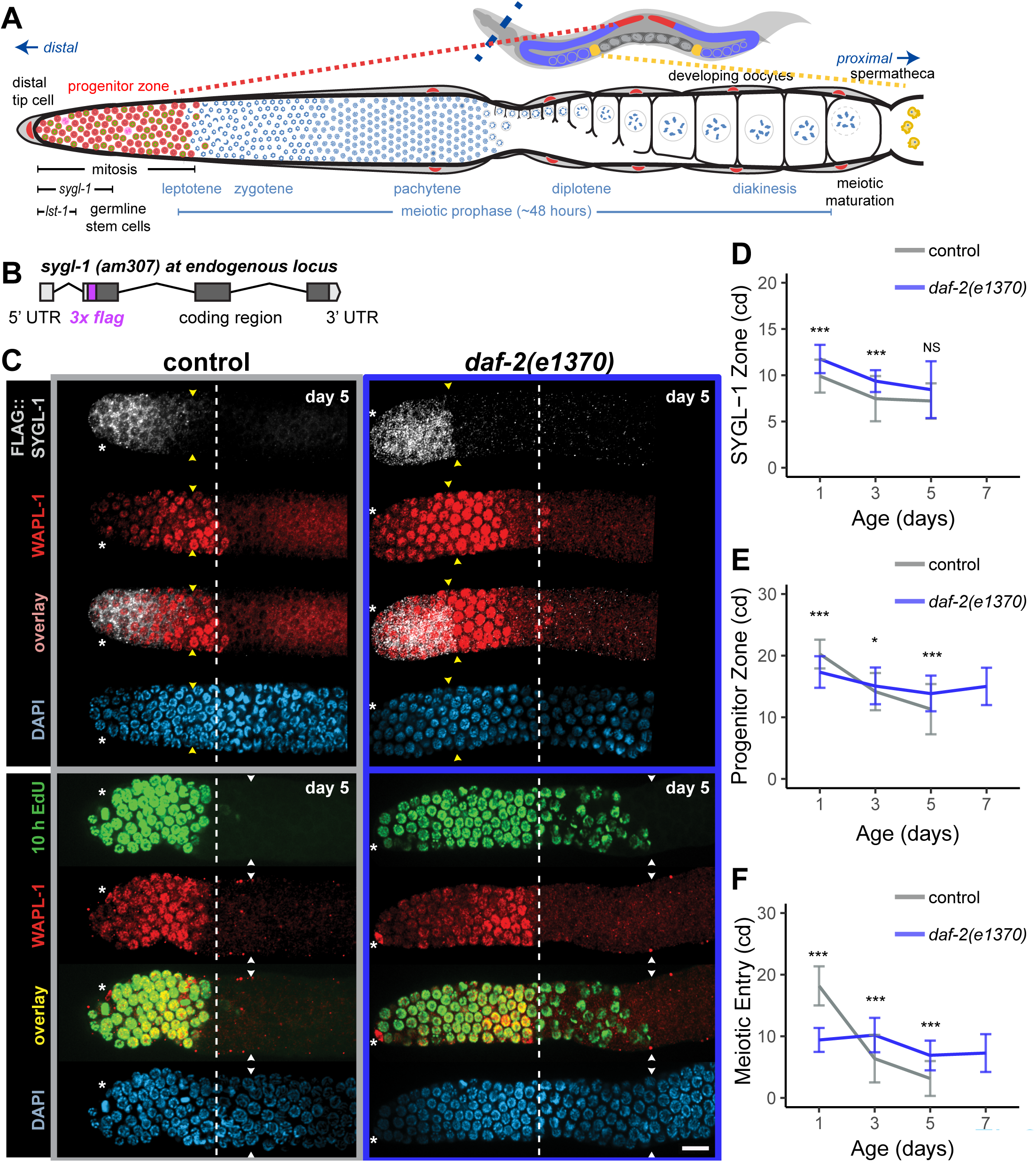
The *daf-2(e1370)* mutation delayed age-related deterioration of germ cells. A) One of two gonad arms of the young adult hermaphrodite. Cells progress from mitotic cycling to meiotic prophase to meiotic maturation before being fertilized by sperm in the spermatheca (yellow). The progenitor zone (red, defined by WAPL-1 staining) contains mitotically cycling stem cells and progenitor cells. The distal tip cell (nucleus in red, as are other somatic gonad cells) provides LAG-2/delta ligand which interacts with GLP-1/Notch receptor on germs cells to maintain the germline stem cell fate via *sygl-1* and lst-1. Panel A modified with permission from Kocsisova et al., (2019). B) Diagram of the *sygl-1*(am307) genomic locus. DNA encoding a 3xFLAG epitope (magenta) was inserted in-frame using CRISPR/Cas9 genome editing, resulting in an N-terminally tagged fusion protein expressed from the endogenous locus. C) Representative confocal micrograph of distal germlines at day 5. The genotypes were: top left control - *sygl-1*(am307flag); bottom left control – wt; top right - *sygl-1*(am307flag); *daf-2(e1370)* ; bottom right - *daf-2(e1370)*. Top: animals with sygl-1*(am307flag)* were stained for FLAG (gray, row 1), WAPL-1 (red, row 2), FLAG and WAPL-1 overlay (row 3), and DAPI (blue, row 4). Bottom: a second set of animals were exposed to EdU for 10 h and stained for EdU (green, row 5), WAPL-1 (red, row 6), EdU and WAPL-1 overlay (row 7), and DAPI (blue, row 8). Asterisks indicate DTC nucleus position; white dashed line indicates proximal boundary of WAPL-1-positive cells; yellow arrowheads indicate proximal boundary of FLAG:SYGL-1; white arrowheads indicate proximal boundary of EdU. Note: all z-planes were checked to determine proximal boundaries. Scale bar 10 μm. D) The extent of SYGL-1 in cell diameters (cd) measured from the distal tip to the last row with half or more SYGL-1 positive cells. The genotypes were: control-*sygl-1*(am307flag), treatment-*sygl-1*(am307flag); daf-2(e1370). E) The length of the progenitor zone in cell diameters (cd), measured from the distal tip to the last row with half or more WAPL-1 positive nuclei. F) The extent of meiotic entry in cell diameters (cd), measured from the end of the progenitor zone to the last EdU+ nucleus following a 10 hour EdU label. D-F) Bar and whiskers indicate mean ± SD. Kruskal–Wallis test. * P<0.05, ** P<.001, *** P<.0001.

To measure the length of the progenitor zone, we used the WAPL-1 antibody and counted the number of cell diameters from the distal tip to the last row with half or more WAPL-1 immunoreactive nuclei (Mohammad et al., 2018). Wild-type hermaphrodites displayed an age-related decrease in the length of the progenitor zone from 20±2 cd at day 1, to 14±3 cd at day 3, to 11±4 cd at day 5 (**Figure 3C, E, Table S3**, Kocsisova et al., 2019). The progenitor zone of *daf-2(e1370)* mated hermaphrodites at day 1 (17±3 cd) was significantly shorter than wild type. This result is similar to the results of Michaelson et al., (2010) and Qin and Hubbard (2015), who report that the progenitor zone of self-fertile young adult *daf-2(e1370)* animals contains fewer nuclei than wild type. The age-related decline of progenitor zone length was blunted in *daf-2(e1370)* mated hermaphrodites; at day 5, *daf-2(e1370)* hermaphrodites displayed a significantly longer progenitor zone (14±3 cd) than wild type (**Figure 3E, Table S3**). Germline deterioration by day 7 in wild-type hermaphrodites was so severe that it was not possible to reliably measure progenitor zone length. By contrast, day 7 *daf-2(e1370)* hermaphrodites could be analyzed and displayed a progenitor zone that was longer than day 5 wild type. Thus, the length of the progenitor zone was well correlated with progeny production.

To measure meiotic entry, we fed hermaphrodites the nucleoside analog EdU for 10 hours and then dissected, fixed, stained for cells that incorporated EdU, and imaged the germline. The number of rows of cells that exhibited EdU signal was counted. Wild-type hermaphrodites displayed an age-related decline in the extent of meiotic entry from 18±3 cd at day 1, to 6±4 cd at day 3, to 3±3 cd at day 5 (**Figure 3C, F, Table S3**). Day 7 wild type were so deteriorated that it was not possible to measure meiotic entry. The meiotic entry of *daf-2(e1370)* hermaphrodites at day 1 (9±2 cd) was significantly shorter than wild type. The age-related decline of meiotic entry was blunted in *daf-2(e1370)* hermaphrodites; at day 3 and 5, *daf-2(e1370)* hermaphrodites displayed significantly higher meiotic entry (10±3 cd and 7±2 cd) compared to wild type. At day 7, the germline of *daf-2(e1370)* hermaphrodites subjectively appeared much healthier and younger than wild type. Objectively, *daf-2(e1370)* hermaphrodites displayed measurable meiotic entry at day 7 (7±3 cd) that was significantly higher than day 5 wild type (**Figure 3F**). To better understand the relationship between progenitor zone length and meiotic entry, we compared the extent of meiotic entry relative to the length of the progenitor zone. We observed a similar pattern of decline, suggesting that the decline in meiotic entry is not fully explained by the shorter progenitor zone. (**Figure S1H, Table S3**). In general, meiotic entry was well correlated with progeny production.

Overall, the *daf-2(e1370)* mutation delayed reproductive aging measured by progeny production late in life, and it also delayed age-related decreases in the extent of the Notch effector SYGL-1, the length of the progenitor zone, and meiotic entry from the progenitor zone. These results are consistent with the hypothesis that the age-related decrease in Notch signaling reduces stem cell number and function, leading to a decrease in meiotic entry and the age-related decrease in progeny production.

### Immune activation and dietary restriction caused by *eat-2(ad465)* and *phm-2(am117)* delayed reproductive aging and age-related changes in germ cells

Mutations in *eat-2* and *phm-2* disrupt the pharynx and allow live E. coli bacteria to enter the intestine, resulting in activation of an immune response, bacterial avoidance behavior, dietary restriction, and extended lifespan (Kumar et al., 2019). In hermaphrodites with sufficient sperm, these mutations delay reproductive aging: the number of progeny produced in late life (after day 10) is significantly higher in mated *eat-2(ad465)* and *phm-2(am117)* hermaphrodites than in mated wild-type hermaphrodites (Hughes et al., 2007, 2011; Kumar et al., 2019). To quantitatively evaluate these mutants, we first analyzed progeny production. Early progeny production by mated *eat-2(ad465)* hermaphrodites (110±40) was significantly lower than wild-type hermaphrodites (400±90), whereas mid-life progeny production was not significantly different than wild type (70±30 versus 100±30) and late progeny production was significantly higher than wild type (65±30 versus 10±10) (**Figure 4A, C, Table S2**). The total brood size of *eat-2(ad465)* hermaphrodites (245±80) was significantly lower than wild-type hermaphrodites (490±120) (**Figure 4D, Table S2**). For mated *phm-2(am117)* hermaphrodites, early (70±30) and mid-life (40±10) progeny production was significantly lower than wild-type hermaphrodites (400±90 and 100±30). Late progeny production was higher than wild type (26±30 versus 10±10), but the difference was not significant with this sample size (P=0.21) (**Figure 4B-C, Table S2**). The total brood size of *phm-2(am117)* hermaphrodites (130±60) was significantly lower than wild-type hermaphrodites (490±120) (**Figure 4D, Table S2**). Thus, *eat-2* and *phm-2* are necessary to promote high levels of early progeny production and rapid reproductive aging.

**Figure 4:**
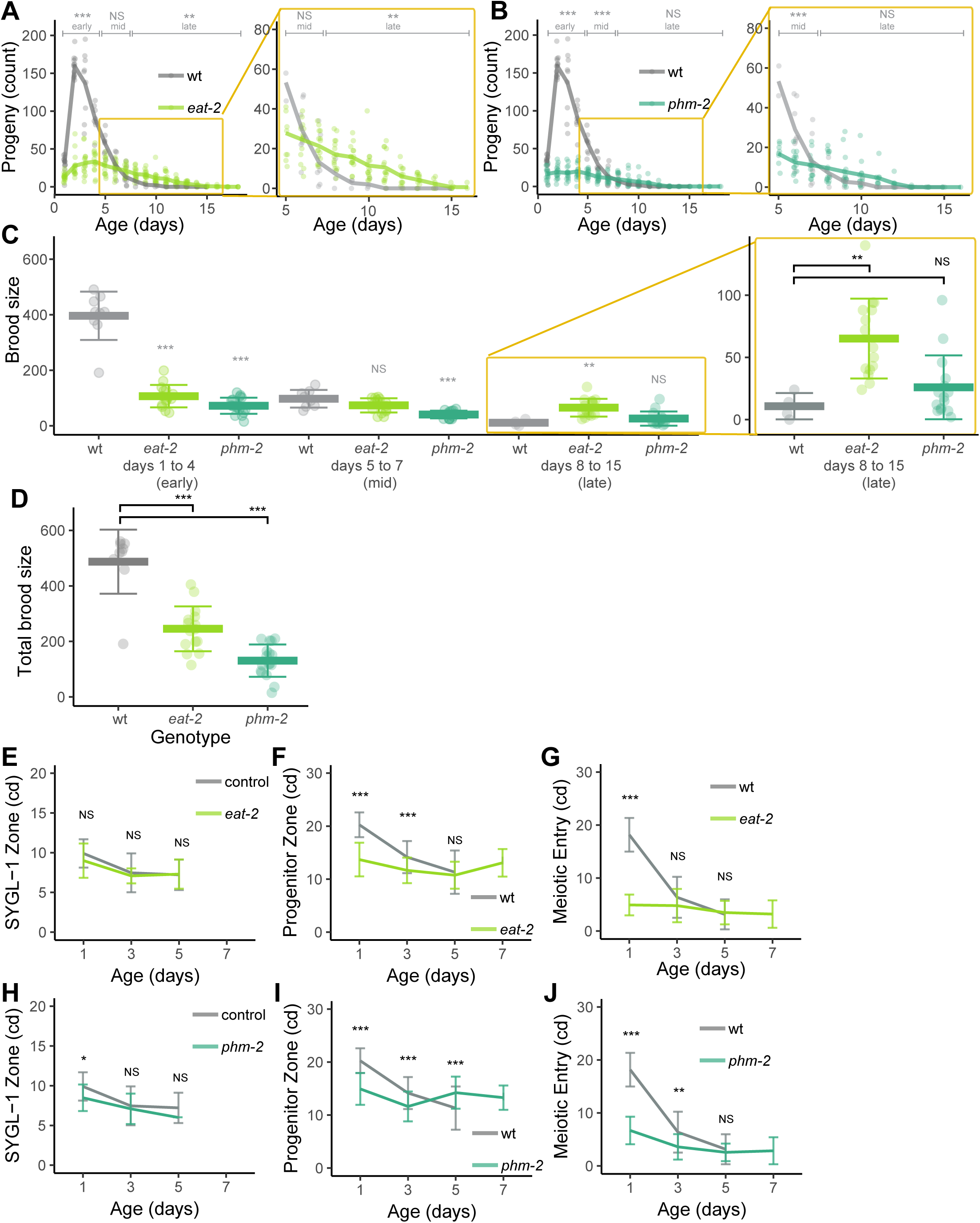
The *eat-2(ad465)* and *phm-2(am117)* mutations delayed reproductive aging and age-related deterioration of germ cells. Wild type (gray), *eat-2(ad465)* (light green), and *phm-2(am117)* (dark green) hermaphrodites were mated to males. A,B) Lines are average daily progeny production, and points are data for individuals. Brackets above indicate early (days 1-4), mid-life (days 5-7) and late (days 8-15) progeny production periods compared by a Wilcoxon Rank Sum test. The area in the yellow box is enlarged on the right. C) The early, mid-life, and late brood size. The area in the yellow box is enlarged on the right. Statistical tests compare wt and mutant at the same days. D) The total brood size, sum of progeny on days 1-15, compared by a Wilcoxon Rank Sum test. E,H) The extent of SYGL-1 in cell diameters (cd) measured from the distal tip to the last row with half or more SYGL-1 positive cells. The genotypes were: control - *sygl-1*(am307flag); *eat-2* - *sygl-1*(am307flag); *eat-2(ad465); phm-2* - sygl-1(am307flag); *phm-2(am117)*. F,I) The length of the progenitor zone in cell diameters (cd), measured from the distal tip to the last row with half or more WAPL-1 positive cells. G,J) The extent of meiotic entry in cell diameters (cd), measured from the end of the progenitor zone to the last EdU+ cell following a 10 hour EdU label. C,E-J) Bar and whiskers indicate mean ± SD. Kruskal–Wallis test. * P<0.05, ** P<.001, *** P<.0001.

To determine the molecular and cellular changes in *eat-2* and *phm-2* hermaphrodites that may be responsible for delayed reproductive aging, we measured germline phenotypes in day 1, 3, 5, and 7 mated hermaphrodites. For *eat-2(ad465);sygl-1(am307flag)* and *phm-2(am117)*;*sygl-1(am307flag)* mated hermaphrodites, the extent of SYGL-1 at all ages was similar or slightly less compared to control (**Figure 4E,H; Table S3**). The progenitor zone of *eat-2(ad465)* mated hermaphrodites at day 1 (14±3 cd) was significantly shorter than that of wild-type mated hermaphrodites (20±2 cd), consistent with reports that in self-fertile young adult *eat-2(ad465)* hermaphrodites the progenitor zone is significantly shorter and contains fewer nuclei than wild type (Brenner and Schedl, 2016; Korta et al., 2012). The age-related decline of progenitor zone length was blunted in *eat-2(ad465)* mated hermaphrodites; at day 5, *eat-2(ad465)* hermaphrodites displayed a progenitor zone (11±3 cd) that was similar to wild type (**Figure 4F,I; Table S3**). Day 7 *eat-2(ad465)* hermaphrodites could be analyzed and displayed a progenitor zone that was longer than day 5 wild type. *phm-2(am117)* mated hermaphrodites displayed a similar pattern – the progenitor zone was shorter than wild type on day 1 and 3, but the age-related decline was blunted, and the progenitor zone was significantly longer than wild type by day 5 (14±3 cd versus 11±4 cd) (**Figure 4F,I; Table S3**). Thus, the length of the progenitor zone was well correlated with progeny production.

The meiotic entry of mated *eat-2(ad465)* hermaphrodites at day 1 (5±2 cd) was significantly lower than wild type (18±3 cd), consistent with a lower level of early progeny production. The age-related decline of meiotic entry was blunted in *eat-2(ad465)* hermaphrodites; at day 3 and 5 the values were similar to wild type, and at day 7 the germline appeared healthier than wild type and displayed measurable meiotic entry (**Figure 4G,J; Table S3**). A similar pattern was observed comparing the extent of meiotic entry relative to the length of the progenitor zone rather than cell diameters (**Figure S2B,E; Table S3**). *phm-2(am117)* mated hermaphrodites displayed a similar pattern. Meiotic entry was significantly lower than wild type in day 1 and 3 hermaphrodites, but the age-related decline was blunted so that the mutants were similar to wild type on day 5 and appeared healthier than wild type and displayed measurable meiotic entry on day 7 (**Figure 4G,J; S2B,E; Table S3**). Thus, meiotic entry was well correlated with progeny production.

Taken together, these results indicate that the pattern of progenitor zone length and meiotic entry matched the pattern of progeny production in *eat-2(ad465)* and *phm-2(am117)* mated hermaphrodites relative to wild type. These results are consistent with the hypothesis that the age-related decrease in progenitor function leads to a decrease in meiotic entry and causes the age-related decrease in progeny production.

### A screen of candidate mutations for delayed reproductive aging identified *che-3(p801)*, which causes defects in sensory perception

To identify mutations that delay reproductive aging without affecting early progeny production, we analyzed candidate mutations. Strains were analyzed because they were reported or suspected to influence lifespan and/or reproduction, and included mutations that affect the TGF-β, DNA damage response, serotonin biosynthesis, mTOR signaling, and sensory perception pathways (**Table S2**, Kocsisova, 2019). Here we focus on the results of only one mutation involved in sensory perception, because it caused a unique pattern of delayed reproductive aging during mid-life.

Mutations that disrupt amphid neuron structure and/or function extend lifespan, indicating that sensory perception of abundant food or some other environmental cue accelerates aging (Alcedo and Kenyon, 2004; Apfeld and Kenyon, 1999). Treatment with the anticonvulsant drug ethosuximide extends lifespan and functions by inhibiting the activity of amphid neurons (Collins et al., 2008a; Evason et al., 2005). Hughes et al. reported that ethosuximide treatment can delay reproductive aging (2007). To investigate the role of sensory perception in reproductive aging, we analyzed nine mutations that impair amphid neuron function affecting five different genes: *che-3(p801), che-3(e1124), che-3(am162), odr-10(ky225), osm-3(p802), osm-3(am172), osm-3(am177), osm-5(p813), and tax-4(p678)* (**Table 1, S2**) (Apfeld and Kenyon, 1999; Collins et al., 2008b; Lans and Jansen, 2007; Mair et al., 2011). An analysis of average progeny number on and after day 8 showed that eight of these mutant strains did not produce significantly more progeny than wild type (range: 10-24 versus 16-24). However, *che-3(p801)* produced 36±16 progeny, which was significantly more than wild-type (22±15).

**Table 1:**
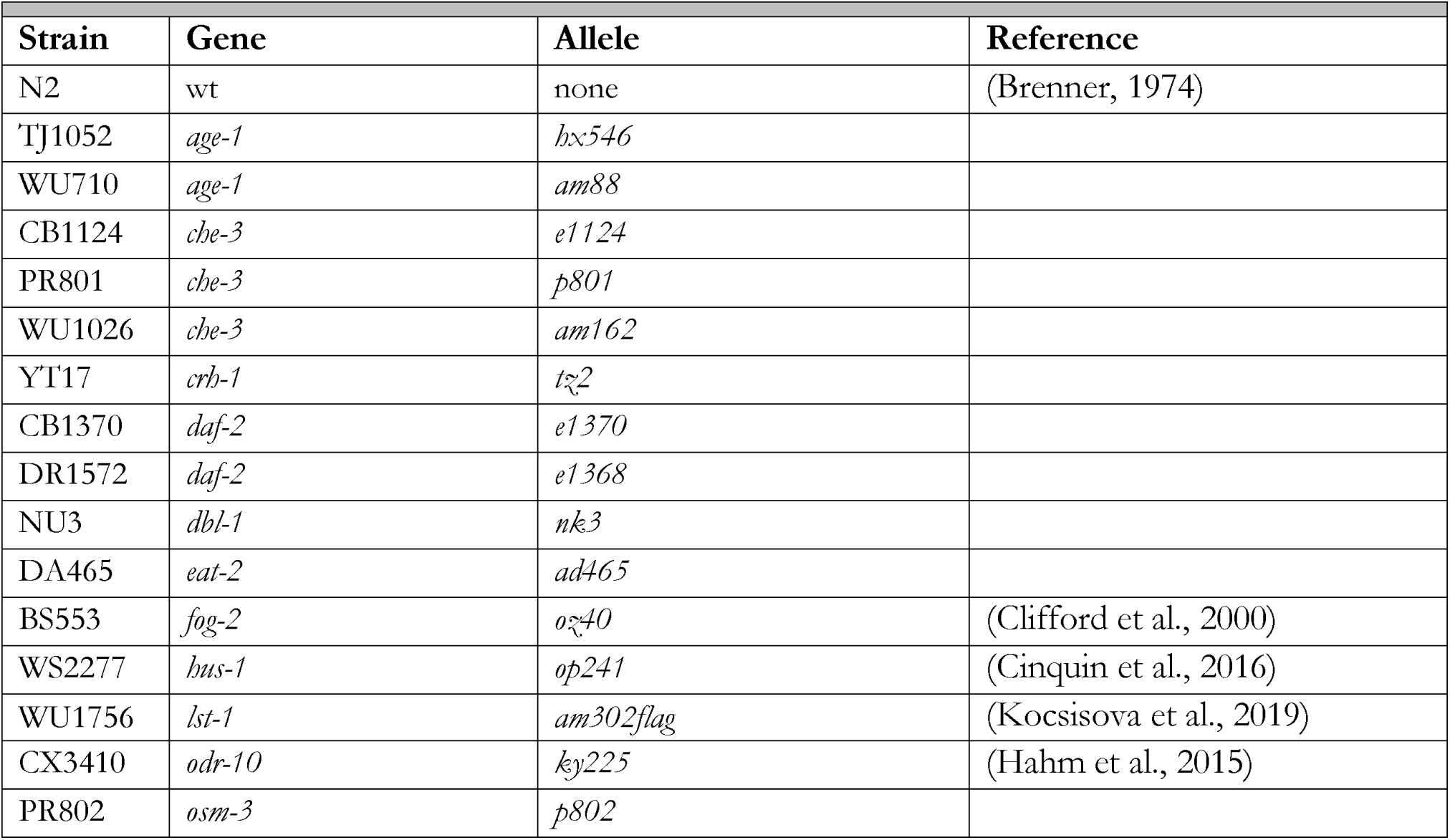
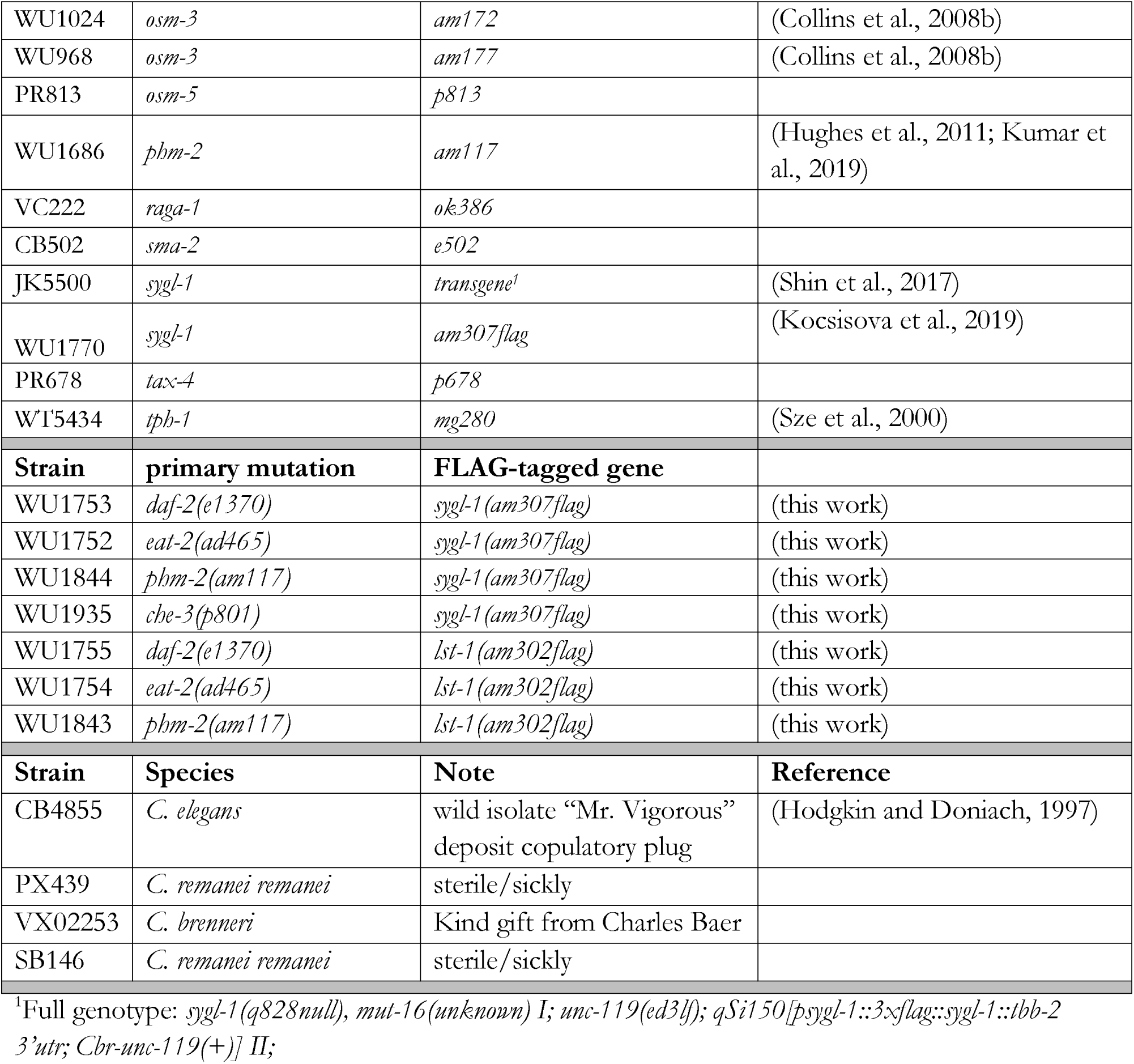
Strains used in this study.

An analysis of complete progeny production by mated *che-3(p801)* hermaphrodites showed that early progeny production (385±40) was similar to wild-type hermaphrodites (380±90), mid-life progeny production was significantly higher than wild type (150±40 versus 90±35), and late progeny production was slightly higher than wild type, although this difference was not significant with this sample size (24±14 versus 16±9) (**Figure 5A-C, Table S2**). The total brood size of *che-3(p801)* hermaphrodites (555±80) was higher than wild-type hermaphrodites (485±120), although this difference was not significant with this sample size (**Table S2**).

**Figure 5:**
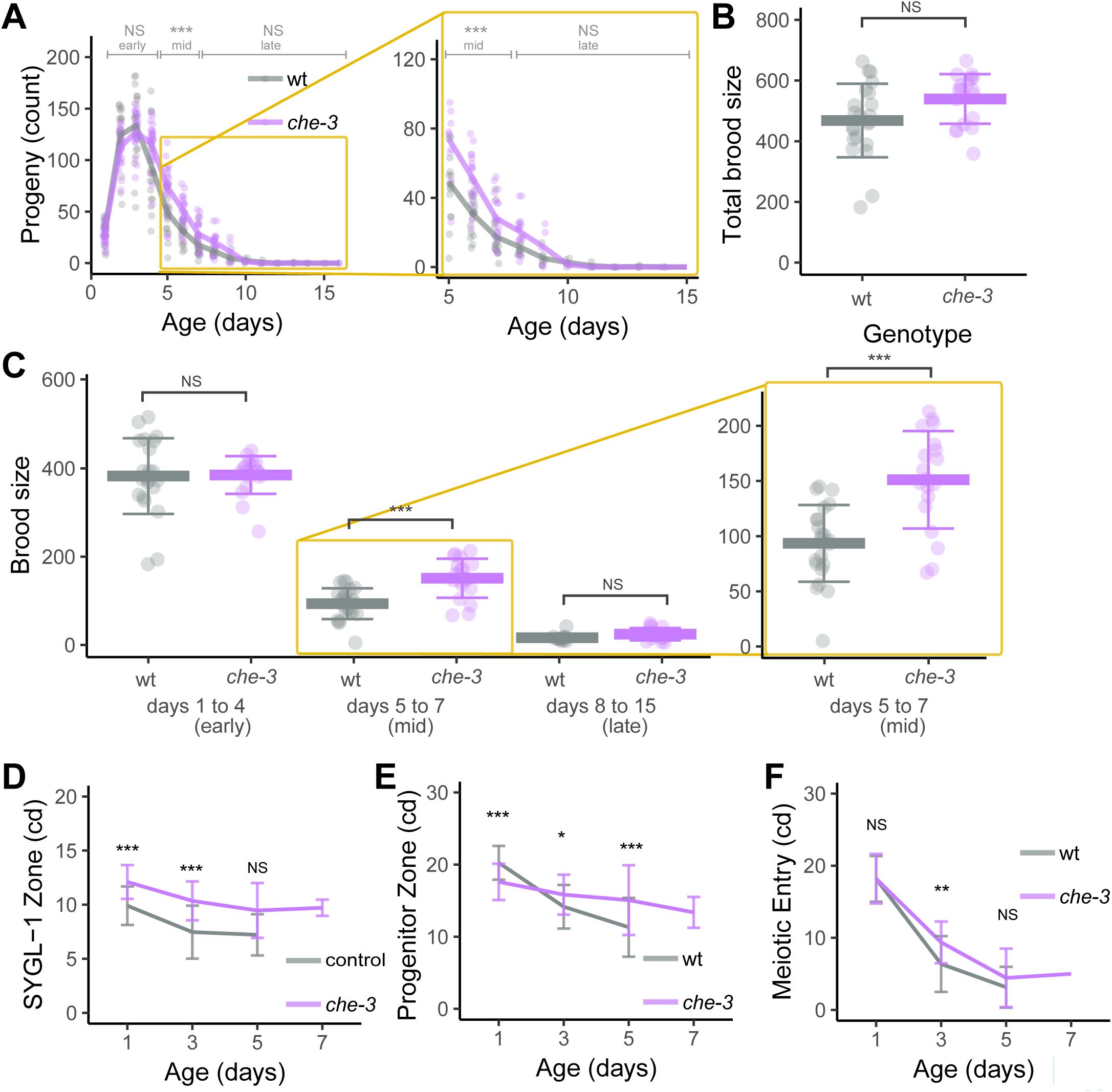
The *che-3(p801)* mutation delayed mid-life reproductive aging and age-related deterioration of germ cells. Wild type (gray) and *che-3(p801)* (purple) hermaphrodites were mated to males. A) Lines are average daily progeny production, and points are data for individuals. Brackets above indicate early (days 1-4), mid-life (days 5-7) and late (days 8-15) progeny production periods compared by a Wilcoxon Rank Sum test. The area in the yellow box is enlarged on the right. B) The total brood size, sum of progeny on days 1-15, compared by a Wilcoxon Rank Sum test. C) The early, mid-life, and late brood size. The area in the yellow box is enlarged on the right. D) The extent of SYGL-1 in cell diameters (cd) measured from the distal tip to the last row with half or more SYGL-1 positive cells. The genotypes were: control - *sygl-1*(am307flag); *che-3* - *sygl-1*(am307flag); che-3(p801). E) The length of the progenitor zone in cell diameters (cd), measured from the distal tip to the last row with half or more WAPL-1 positive cells. F) The extent of meiotic entry in cell diameters (cd), measured from the end of the progenitor zone to the last EdU+ cell following a 10 hour EdU label. C-F) Bar and whiskers indicate mean ± SD. Kruskal–Wallis test. * P<0.05, ** P<.001, *** P<.0001.

Thus, *che-3(p801)* displayed a different pattern of delayed reproductive aging compared to *daf-2(e1370), eat-2(ad465)*, and *phm-2(am117)*. Mated *che-3(p801)* hermaphrodites produced wild-type levels of progeny on days 1-4, significantly exceeded wild-type levels on days 5-7, and in some replicates exceeded wild-type levels on day 8 and beyond. Notably, *che-3* mutants outperformed wild-type animals in middle and old age without a decrease in peak progeny production in young animals.

### Impaired sensory perception caused by *che-3(p801)* delayed mid-life reproductive aging and age-related changes in germ cells

To determine the molecular and cellular changes in *che-3* hermaphrodites that may be responsible for delayed reproductive aging, we measured germline phenotypes in day 1, 3, 5, and 7 mated hermaphrodites. Mated che-3(p801);*sygl-1(am307flag)* hermaphrodites displayed significantly longer SYGL-1 zone at days 1 and 3 compared to control (12±2 versus 10±2 cd) and (10±2 versus 7.5±2 cd) (**Figure 5D, Table S3**). This trend continued on day 5, although the difference was not significant with both statistical tests (9.5±2.5 versus 7±2 cd). The SYGL-1 extent of che-3(p801);sygl-1*(am307flag)* hermaphrodites was measurable on day 7 (10±1 cd), when wild type is too degenerated to analyze, and it was greater than the wild type value on day 5.

The progenitor zone of *che-3(p801)* mated hermaphrodites at day 1 (18±2.5cd) was significantly shorter than that of wild-type mated hermaphrodites (20±2 cd). The age-related decline of progenitor zone length was blunted in *che-3(p801)* mated hermaphrodites; at days 3 and 5, *che-3(p801)* hermaphrodites displayed a progenitor zone that was significantly longer than wild type (16±3 versus 14±3 cd; 15±5 versus 11±4 cd) (**Figure 5E; Table S3**). Day 7 *che-3(p801)* hermaphrodites could be analyzed and displayed a progenitor zone that was longer than day 5 wild type. Thus, the length of the progenitor zone was well correlated with progeny production.

The meiotic entry of mated *che-3(p801)* hermaphrodites at day 1 (18±3 cd) was similar to wild type (18±3 cd), consistent with a similar level of early progeny production. The age-related decline of meiotic entry was blunted in *che-3(p801)* hermaphrodites; at day 3 the value was significantly higher than wild type (9±3 versus 6±4 cd), and at day 5 the value was slightly higher than wild type, although this difference was not significant with this sample size (4±4 versus 3±3 cd) (**Figure 5F; Table S3**). A similar pattern was observed comparing the extent of meiotic entry relative to the length of the progenitor zone rather than cell diameters: *che-3(p801)* hermaphrodites displayed significantly higher meiotic entry compared to wild type at days 1 and 3 (**Figure S3B; Table S3**). Thus, meiotic entry was well correlated with progeny production.

Thus, the *che-3(p801)* mutation that increased the number of progeny produced in mid-life also increased extent of the Notch effector SYGL-1, the length of the progenitor zone, and meiotic entry from the progenitor zone in mid-life. These results support the hypothesis that the age-related decrease in progenitor function leads to a decrease in meiotic entry and causes the age-related decrease in progeny production.

### Ectopic expression of the Notch pathway effector SYGL-1 was sufficient to delay mid-life reproductive aging and age-related changes in germ cells

The distal tip cell expresses the Notch ligand LAG-2, which binds GLP-1 Notch expressed by the germline stem cells, thereby maintaining these cells in the proliferative, stem cell fate (Austin and Kimble, 1987; Kimble and Ward, 1988). Kocsisova et al., (2019) reported an age-related decline in the length and size of the Notch signaling domain, as measured by the expression of the direct Notch target genes lst-1 and *sygl-1*. Based on these observations, we hypothesized that an age-related decrease in Notch pathway signaling causes the decrease in stem cells, which leads to age-related decreases in progenitor zone output and progeny production. This hypothesis predicts that sustaining the activity of the Notch pathway in aging animals could delay reproductive aging.

Shin et al. (2017) showed that the 3’ untranslated region (UTR) of the *sygl-1* mRNA is a site of negative regulation. Transgenic animals in which the *sygl-1* 3’UTR is replaced by the unregulated tbb-2 3’UTR result in ectopic expression of SYGL-1 in the distal germline, expanding the zone of accumulation proximally (Shin et al., 2017). We refer to this ectopic expression strain as *sygl-1*(ee) (**Figure 6A**).

**Figure 6:**
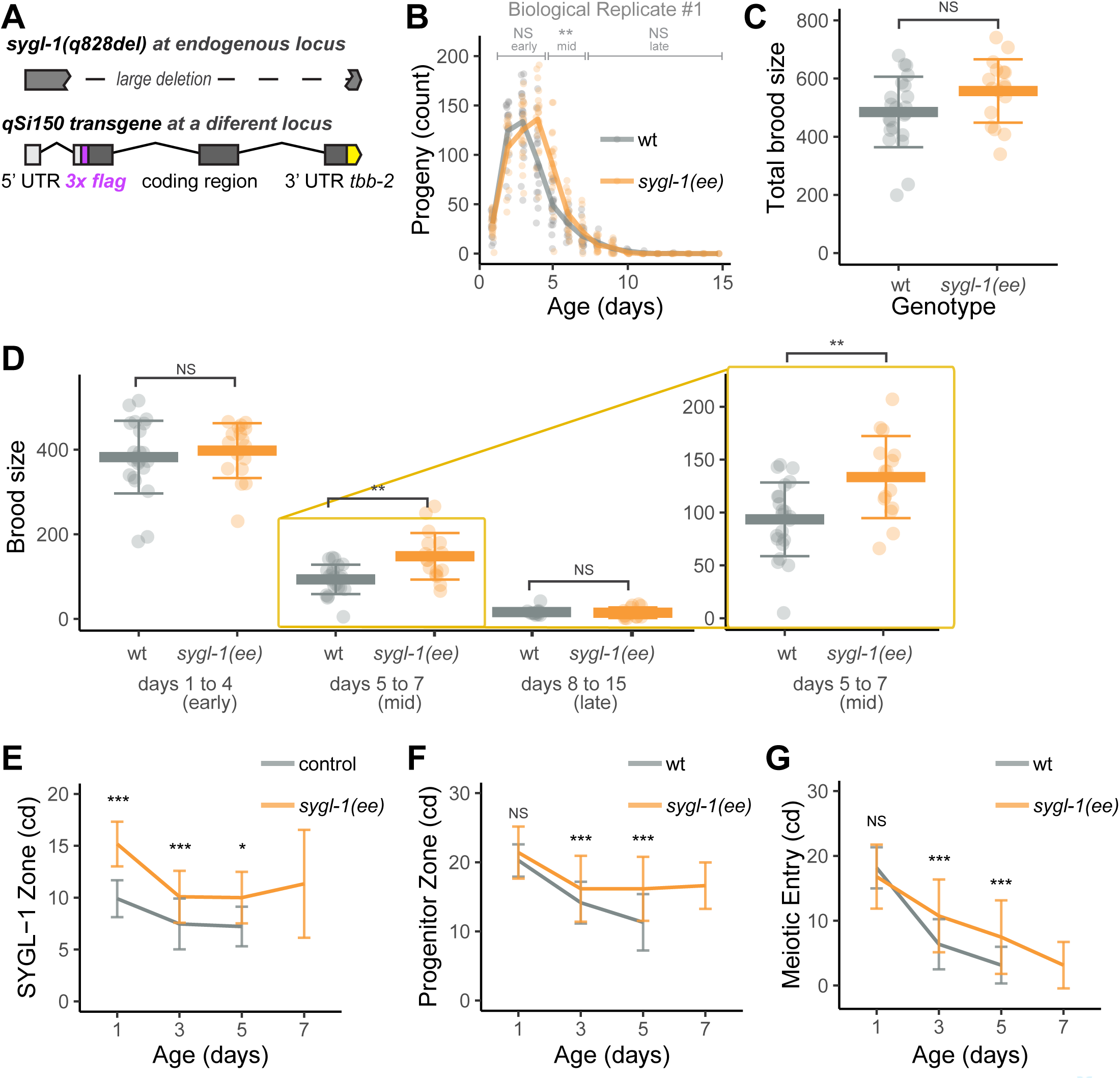
Ectopic expression of SYGL-1 delayed mid-life reproductive aging and age-related deterioration of germ cells. Wild type (gray) and *sygl-1*(ee) (orange) hermaphrodites were mated to males. A) Genotype of the sygl-1(ee) strain (JK5500) is *sygl-1*(q828null), mut-16(unknown) I; unc-119(ed3lf); qSi150[p*sygl-1*::3xflag::sygl-1::tbb-2 3’utr; Cbr-unc-119(+)] II. Diagrams show the endogenous *sygl-1* locus containing the q828 deletion mutation (top) and the integrated transgene containing the *sygl-1* promoter, an in-frame 3X FLAG epitope (magenta), and the 3’ UTR from tbb-2 (yellow) (bottom) (Shin et al., 2017). The sygl-1(ee) strain contains the q828 deletion at the endogenous locus and the qSi150 transgene integrated at another location in the genome. B) Lines are average daily progeny production, and points are data for individuals. Brackets above indicate early (days 1-4), mid-life (days 5-7) and late (days 8-15) progeny production periods compared by a Wilcoxon Rank Sum test. C) The total brood size, sum of progeny on days 1-15, compared by a Wilcoxon Rank Sum test. D) The early, mid-life, and late brood size. The area in the yellow box is enlarged on the right. E) The extent of SYGL-1 in cell diameters (cd) measured from the distal tip to the last row with half or more SYGL-1 positive cells. The genotypes were: control - *sygl-1*(am307flag); *sygl-1*(ee) - *sygl-1*(q828null), mut-16(unknown) I; unc-119(ed3lf); qSi150[p*sygl-1*::3xflag::*sygl-1*::tbb-2 3’utr; Cbr-unc-119(+)]. F) The length of the progenitor zone in cell diameters (cd), measured from the distal tip to the last row with half or more WAPL-1 positive cells. G) The extent of meiotic entry in cell diameters (cd), measured from the end of the progenitor zone to the last EdU+ cell following a 10 hour EdU label. C-G) Bar and whiskers indicate mean ± SD. Kruskal–Wallis test. * P<0.05, ** P<.001, *** P<.0001.

To determine if the expanded accumulation of SYGL-1 is sustained, we used anti-FLAG antibodies to measure the domain of SYGL-1. Mated *sygl-1*(ee) hermaphrodites displayed significantly longer SYGL-1 extents at days 1, 3, and 5 compared to wild-type (15±3 versus 10±2 cd; 10±2.5 versus 7.5±2 cd; 10±2.5 versus 7±2 cd, respectively.) (**Figure 6E, Table S3**). Day 7 *sygl-1*(ee) hermaphrodites displayed measurable SYGL-1 expression (11±5 cd), when wild type is too degenerated to analyze. Thus, mutating the 3’UTR of the *sygl-1* transgene resulted in expanded accumulation of SYGL-1 protein that was sustained into mid-life.

To determine if expanded accumulation of SYGL-1 is sufficient to delay reproductive aging, we measured daily progeny production of mated *sygl-1*(ee) hermaphrodites in two biological replicates. Early progeny production was slightly but not significantly higher than wild type in both replicates (400±65 versus 380±90, and 530±40 versus 510±30). Mid-life progeny production was significantly higher than wild type in both replicates (150±55 versus 90±35, and 180±60 versus 100±40). Late progeny production was not significantly different than wild type (15±13 versus 16±9, and 23±22 versus 9±7) (**Figure 6B-D, S4B-E, Table S2**). The total brood size of *sygl-1*(ee) hermaphrodites was higher than wild-type hermaphrodites (560±110 versus 485±120, and 730±90 versus 610±60), and this difference was significant in the second replicate (**Table S2**). Thus, *sygl-1*(ee) hermaphrodites displayed a pattern of delayed reproductive aging similar to *che-3(p801)* hermaphrodites, producing wild-type levels of progeny on days 1-4 and significantly exceeding wild-type levels on days 5-7. Notably, *sygl-1*(ee) mutants outperformed wild-type animals in middle age without a decrease in peak progeny production in young animals.

To determine the cellular changes in *sygl-1*(ee) hermaphrodites that may be responsible for delayed reproductive aging, we measured germline phenotypes in day 1, 3, 5, and 7 mated hermaphrodites. The progenitor zone of *sygl-1*(ee) mated hermaphrodites at day 1 (21±4 cd) was slightly but not significantly longer than wild-type mated hermaphrodites (20±2 cd). The age-related decline of progenitor zone length was blunted in *sygl-1*(ee) mated hermaphrodites; at days 3 and 5, *sygl-1*(ee) hermaphrodites displayed a progenitor zone that was significantly longer than wild type (16.5±5 versus 14±3 cd; 16±5 versus 11±4 cd) (**Figure 6F; Table S3**). Day 7 *sygl-1*(ee) hermaphrodites could be analyzed and displayed a progenitor zone that was longer than day 5 wild type. Thus, the length of the progenitor zone was well correlated with progeny production.

The meiotic entry of mated *sygl-1*(ee) hermaphrodites at day 1 (17±5 cd) was similar to wild type (18±3 cd), consistent with a similar level of early progeny production. The age-related decline of meiotic entry was blunted in *sygl-1*(ee) hermaphrodites; at days 3 and 5, the values were significantly higher than wild type (11±6 versus 6±4 cd, and 7.5±6 versus 3±3 cd) (**Figure 6G; Table S3**). A similar pattern was observed comparing the extent of meiotic entry relative to the length of the progenitor zone rather than cell diameters - *sygl-1*(ee) hermaphrodites displayed significantly higher meiotic entry compared to wild type at days 3 and 5 (**Figure S4G; Table S3**). Thus, meiotic entry was well correlated with progeny production.

The *sygl-1*(ee) transgene that increased the number of progeny produced in mid-life also increased the length of the progenitor zone and meiotic entry from the progenitor zone. These results support the hypothesis that the age-related decrease in progenitor function leads to a decrease in meiotic entry and causes the age-related decrease in progeny production.

To determine if ectopic expression of SYGL-1 in the germline affected somatic aging, we measured the lifespan of unmated *sygl-1*(ee) hermaphrodites. These animals displayed a mean lifespan that was not significantly different from wild-type (WT: 12.5 days; *sygl-1*(ee) 11.9 days; log-rank test P=0.7343) (**Figure S4A**). Thus, ectopic expression of SYGL-1 can uncouple reproductive and somatic aging.

## Discussion

### Post-reproductive lifespan is a feature of both hermaphrodite/male and female/male Caenorhabditids

Of 63 characterized species in the genus *Caenorhabditis*, 60 are female/male whereas only three are hermaphrodite/male. However, these three - *C. elegans, C. briggsae*, and *C. tropicalis* - have been analyzed extensively because they are convenient for laboratory genetic studies (Kiontke et al., 2011). Protandry likely evolved independently three times in the genus *Caenorhabditis*, and the evolution of a radically different mode of reproduction might have affected reproductive aging. Some genes regulating the development of protandry, such as fog-2 in *C. elegans*, are new and unique to the species (Nayak et al., 2005; Schedl and Kimble, 1988). Also, the near-lack of males, small size of self-sperm, radically different ascaroside signaling, and frequency of sperm-depletion might have created new evolutionary pressures on the aging reproductive tract (Angeles-Albores et al., 2017). To evaluate the generality of conclusions based on *C. elegans* self-fertile hermaphrodites, we analyzed two female/male species: *C. brenneri* and *C. remanei*.

Fertility studies of these female/male species are challenging due to inbreeding depression. Strains suitable for laboratory study can be generated through careful mating protocols to achieve a relatively inbred strain that harbors relatively few deleterious, sterile, and lethal alleles. We analyzed such a strain of *C. brenneri* (VX02253, a kind gift from Charles Baer). In the case of *C. remanei*, we obtained the strain from the CGC. In both species, a decline in reproductive function occurred several days before mated females began to die of old age. This is the same pattern previously reported (and confirmed here) in both wild-type protandrous and feminized *C. elegans* (Hodgkin and Barnes, 1991; Hughes et al., 2007). Thus, a post-reproductive lifespan is not likely to be a consequence of an evolutionary history of self-fertile hermaphrodites. This result is consistent with recent findings in another female/male species, C. inopinata, the sister species to *C. elegans* (referred to as C. sp. 34 prior to 2017) (Kanzaki et al., 2018; Woodruff and Phillips, 2018). C. inopinata produce the majority of progeny at days 2-3 of adulthood, few progeny at days 5-7, and are nearly sterile thereafter, while survival remains over 75% until ∼day 15 of adulthood (Woodruff et al., 2019).

Examining reproductive aging in more prevalent female/male species is important for understanding the evolution of reproductive aging. Aging varies across the tree of life (Jones et al., 2014), yet in almost all species that age, reproduction fails earlier in life than somatic functions (Mitteldorf and Goodnight, 2013). It has been hypothesized that the evolution of sperm-limitation as a consequence of protandrous hermaphroditism could have led to the emergence of a post-reproductive lifespan and an adaptive death (Lohr et al., 2019). The results presented here and in Woodruff et al. (2019) suggest the explanation for the post-reproductive lifespan needs to account for both female/male and hermaphrodite/male species.

### Distal germline morphology and stem cell activity were correlated with progeny production in a variety of mutants

Kocsisova et al. (2019) described molecular and cellular age-related changes in the wild-type germline and gonad, leading to the hypothesis that these declines cause reproductive aging. However, the observational studies of Kocsisova et al., (2019) did not rigorously establish the causal role of the reported age-related declines. To address the functional significance of the age-related declines observed in wild-type animals, we measured the same phenotypes in three mutant strains previously reported to display extended reproductive spans and increased numbers of progeny produced in late-life: daf-2(e1370), *eat-2(ad465)*, and *phm-2(am117)* (**Figure 7A**).

**Figure 7:**
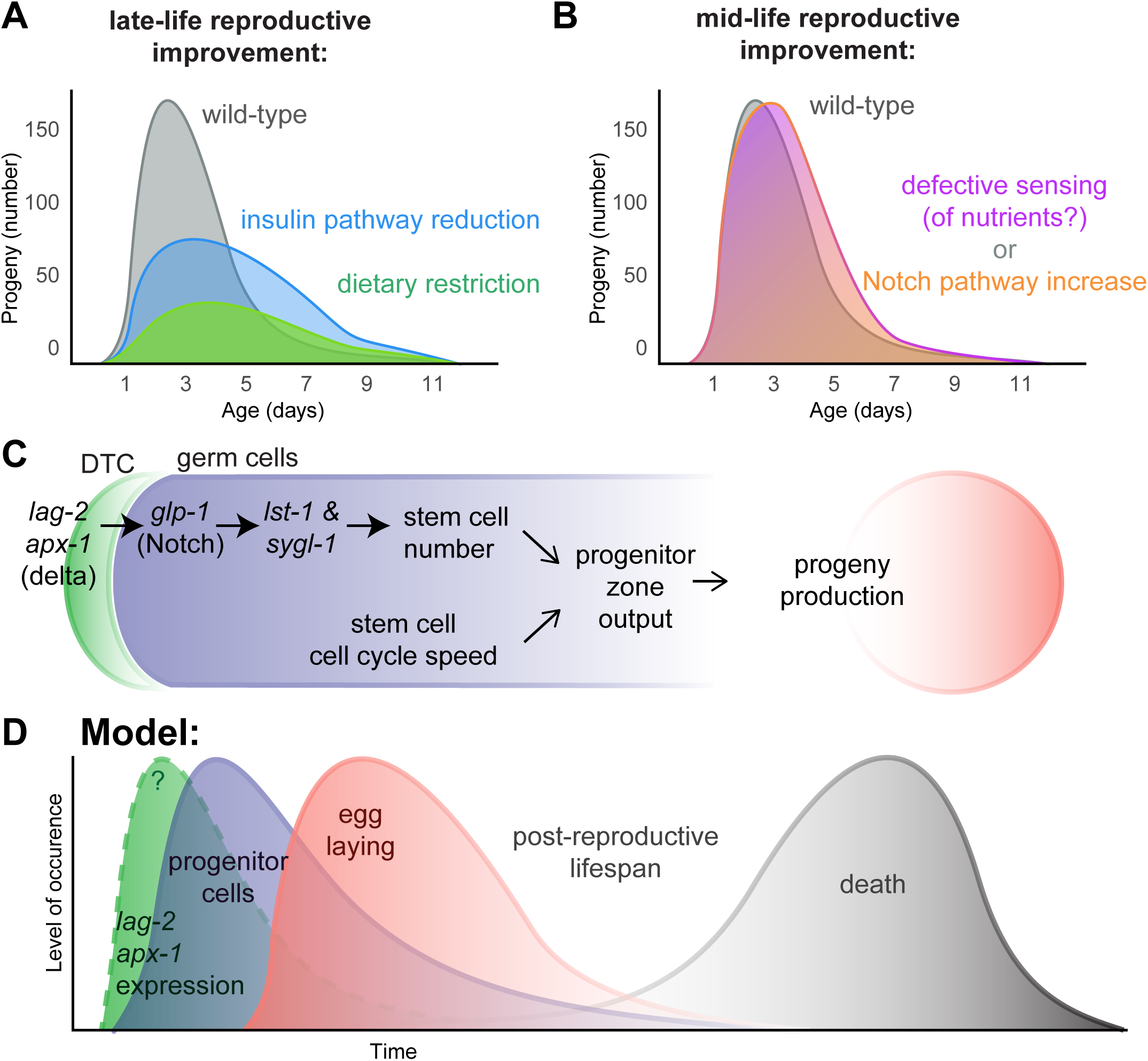
Mid-life reproductive improvement results from loss-of-function in nutrient sensing or gain-of-function in Notch signaling. 6. The model summarizes daily progeny production in mated daf-2(e1370), *eat-2(ad465)*, and phm-2(am117) hermaphrodites that display a decrease in peak progeny production and an increase in late-life reproduction. This pattern is gene and allele specific for the insulin/IGF-1 pathway, since it is displayed by *daf-2(e1370)* but not several other daf-2 or age-1 mutations (Hughes (2005) and this study). The interpretation of *eat-2(ad465)* and *phm-2(am117)* phenotypes are complicated by the finding that these strains are infected by E. coli OP50, resulting in an immune response, bacterial avoidance behavior, and dietary restriction (Kumar et al., 2019). B) The model summarizes daily progeny production in mated *che-3(p801)* and *sygl-1*(ee) hermaphrodites that display normal peak progeny production and increased mid-life reproduction. C) The diagram shows signaling events that promote progeny production. LAG-2 and APX-1 ligand on the distal tip cell, activation of GLP-1 Notch on germ cells, and transcription of target genes lst-1 and *sygl-1* regulate the number of stem cells. The speed of the cell cycle is regulated by unknown factors not included in this diagram. Together, the number of stem cells and cell cycle speed determine progenitor zone output and progeny production. D) The diagram shows an integrated time course linking development and aging. We propose time-limited high LAG-2 and APX-1 expression results in a time-limited pulse of progenitor cells that results in a time-limited pulse of egg laying. A post-reproductive period precedes the pulse of senescent death.

In day 1 adults, the germline of these three mutant animals is smaller, and meiotic entry is lower than in wild-type. Thus, these genes function early in life to promote the establishment of a germline with normal size and function. However, the analysis of day 3, 5, and 7 adults revealed that the rate of decline is more gradual in these mutants compared to wild type. At days 5 and 7, the mutant animals appear more youthful than wild type at days 5 and 7. This effect is most pronounced in daf-2(e1370). The mated *daf-2(e1370)* hermaphrodites began adulthood with a shorter SYGL-1 zone, shorter progenitor zone, and lower rate of meiotic entry than wild-type, and also produced fewer progeny during the first four days of adulthood than wild-type. With age, the decline in mated *daf-2(e1370)* hermaphrodites was less steep than in mated wild-type hermaphrodites. At days 3 and 5 of adulthood, the mated *daf-2(e1370)* hermaphrodites maintained a longer SYGL-1 zone, a longer progenitor zone, and a higher rate of meiotic entry than wild-type, and also produced more progeny in late life than wild-type. Similarly, the patterns of germline morphology and stem cell activity in the other mutant strains matched the patterns of progeny decline. These correlations support the hypothesis that an age-related decrease in progenitor cell function leads to a decrease in meiotic entry and causes the age-related decrease in progeny production. Because correlations do not definitively establish cause, it remains possible that another cause exists that was not measured.

To elucidate the role of the insulin/IGF-1 pathway in reproductive aging, we analyzed several alleles of age-1 and daf-2. The numerous mutations in daf-2 have been described in an allelic series and sorted into two classes: Class I alleles, including e1368 and m41, affect certain extracellular regions of the receptor while Class II alleles, including e1370, are pleiotropic and affect either the ligand binding pocket or the tyrosine kinase domain. Class I and Class II alleles differ in movement, fecundity, and aging phenotypes (Arantes-Oliviera et al., 2002; Ewald et al., 2015; Gems et al., 1998; Patel et al., 2008; Podshivalova et al., 2017). Of the insulin signaling pathway mutants tested in this study and by Hughes (2005) (daf-2(e1368), daf-2(m41), age-1(am88), and age-1(hx546)), only *daf-2(e1370)* extended reproduction and significantly increased progeny production on and after day 8. This result suggests that the e1370 allele creates levels of insulin signaling that are “just-right” to delay reproductive aging without excessively hindering development. However, *daf-2(e1370)* does significantly decrease early reproduction (Hughes, 2005; Hughes et al., 2007; this study).

### Defects in sensory perception lead to mid-life reproductive improvement

Dozens of mutations have been reported to delay somatic aging and extend lifespan, but only a handful have been reported to delay reproductive aging (Scharf et al., 2021). Hughes et al. (2007) analyzed daily progeny production of long-lived mutants; whereas some displayed increased late-life progeny production, such as *daf-2(e1370)* and *eat-2(ad465)*, others did not, such as clk-1(qm30) and isp-1(qm150). Two candidate gene studies reported extended reproductive span in mated hermaphrodites: in the TGF-β pathway by sma-2(e502) (Luo et al., 2010) and in the sodium homeostasis pathway by RNAi-mediated knockdown of nhx-2 and sgk-1(Wang et al., 2014). A forward-genetic screen for extended reproduction by mated hermaphrodites identified that phm-2(am117) increased the number of progeny produced on and after day 10 of adulthood and extended lifespan (Hughes et al., 2011; Kumar et al., 2019). The mechanism of lifespan extension and increased late-life progeny production in both *eat-2(ad465)* and *phm-2(am117)* includes an immune and behavioral response that leads to dietary restriction (Kumar et al., 2019).

To identify new genes involved in this process, we analyzed candidate mutations. The *che-3(p801)* mutant displayed a distinctive pattern of delayed reproductive aging – increased progeny production during midlife, from days 5-7 (**Figure 7B**). It was also unusual because it exhibited wild-type levels of progeny production on days 1-4. Previously reported long-lived mutants displayed decreased early peak progeny production and a decreased total brood size in mated hermaphrodites (Hughes et al., 2007, 2011; Kumar et al., 2019; Wang et al., 2014). The lifespan extension of *che-3(p801)* is caused by defective chemosensation resulting from defects in the amphid neurons (Apfeld and Kenyon, 1999; Collins et al., 2008b). We interpret the pattern of progeny production in *che-3(p801)* to be a result of a particular deficiency in perception, possibly a perception of nutrient deficiency without actual nutrient deficiency. Thus, it is distinct from the immune activation and dietary restriction that likely result in decreased peak progeny production in *eat-2(ad465)* and phm-2(am117). Whereas tradeoff theories propose that reduced progeny production is the cause of extended lifespan, the analysis of *che-3(p801)* demonstrates that it is possible for a mutation to extend life without reducing early progeny production or total brood size (**Figure 7B**).

### Increased Notch signaling leads to mid-life reproductive improvement

Kocsisova et al., (2019) hypothesized that the age-related decline in stem cell number and slowing of the stem cell cycle cause reproductive aging. This hypothesis predicts that increasing the number of stem cells and/or the rate of the stem cell cycle will delay reproductive decline. The pathway that regulates the rate of the stem cell cycle has not been defined, but the Notch signaling pathway is well-established to affect stem cell number (Fox and Schedl, 2015; Lee et al., 2016; Shin et al., 2017). A transgenic strain in which 3’UTR-mediated repression of the Notch effector *sygl-1* is interrupted displays a larger stem cell pool in young adults (Shin et al., 2017). To test whether extending Notch signaling output would be sufficient to delay reproductive aging, we analyzed this *sygl-1*(ee) strain.

Here we report that ectopic expression of *sygl-1* was sufficient to increase progeny production in mid-life (days 5-7), similar to che-3(p801). Unlike previously reported mutants, which increased late-life reproduction and decreased peak reproduction, *sygl-1*(ee) exhibited wild-type levels of reproduction on days 1-4 (**Figure 7B**). Unlike *che-3(p801)* and all other mutant strains reported to delay reproductive aging, this strain was not long-lived. These findings directly support the model that age-related declines in stem cell number contribute to reproductive aging (**Figure 7C**). Based on these findings, we speculate that age-related decreases in the Notch ligand, receptor signaling, or SYGL-1 effector are an underlying cause of reproductive aging (**Figure 7D**).

### Summary and Conclusion

We identified two genetic manipulations that increase mid-life reproduction without decreasing early progeny production: *che-3(p801)* and *sygl-1*(ee). *che-3(p801)* also extends lifespan, which breaks the correlation between reduced progeny production and extended lifespan displayed by *daf-2, eat-2* and *phm-2* mutants. *sygl-1*(ee) delays reproductive aging without extending lifespan, which breaks the correlation between reproductive and somatic aging. This pattern is complementary to many mutations that can extend lifespan but do not delay reproductive aging. These results show that while reproductive and somatic aging are connected, they are also separable. An important difference between nematode and mammalian female germline development is that while mammalian female germline stem cells appear to cease dividing by mitosis before birth, nematode germline stem cell divisions continue into adulthood. Although the results presented here apply to *Caenorhabditis* nematodes and not humans, an analogy to human female reproductive aging may be illustrative: rather than identifying ways to help women have babies in their early 60s (reproductive span extension), we identified processes that are analogous to help women have babies in their late 30s (mid-life reproductive increase), without compromising fertility in their 20s. In *C. elegans* it is possible to increase mid-life reproduction without compromising early/peak reproduction.

## Figures and Tables

**Figure S1:**
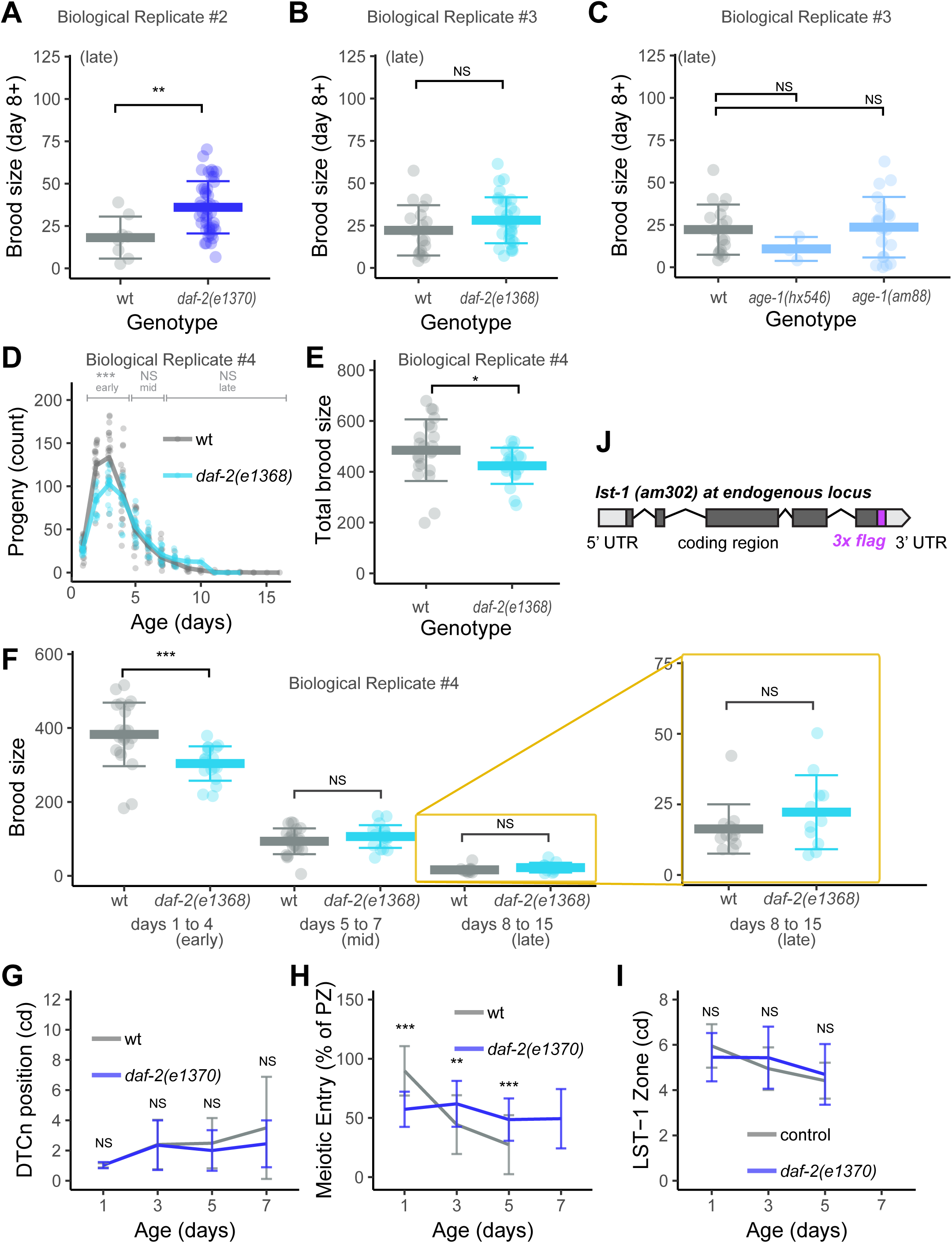
Reproductive and germline phenotypes caused by daf-2 and age-1 mutations. Wild type (gray), *daf-2(e1370)* (dark blue), daf-2(e1368) (turquoise), age-1(hx546) and age-1(am88) (light blue) hermaphrodites were mated to males. A-C) Late brood size is the sum of progeny produced on day 8, 9, 10… until reproductive cessation (up to day 15). Bar and whiskers indicate mean ± SD with individual data points shown. Mutants were compared to a concurrent wild-type control (displayed for each biological replicate) with a Wilcoxon Rank Sum test. D) Lines are average daily progeny production, and points are data for individuals. Brackets above indicate early (days 1-4), mid-life (days 5-7) and late (days 8-15) progeny production periods compared by a Wilcoxon Rank Sum test. E) The total brood size, sum of progeny on days 1-15, compared by a Wilcoxon Rank Sum test. F) The early, mid-life, and late brood size. Area in yellow box is enlarged on the right. G) Values are the distance of the distal tip cell nucleus (DTCn) from the distal tip measured in cell diameters (cd). H) The extent of meiotic entry (measured in cell diameters) expressed as a percent of the extent of the progenitor zone (measured in cell diameters). I) The extent of LST-1 in cell diameters (cd) measured from the distal tip to the last row with half or more LST-1 positive cells. The genotypes were: control – lst-1(am302flag); *daf-2(e1370)* - lst-1(am302flag); *daf-2(e1370)* A-I) Kruskal–Wallis test. * P<0.05, ** P<.001, *** P<.0001. J) Diagram of the lst-1(am302) genomic locus. DNA encoding a 3xFLAG epitope (magenta) was inserted in-frame using CRISPR/Cas9 genome editing, resulting in a C-terminally tagged fusion protein expressed from the endogenous locus. We chose a C-terminal fusion to achieve labeling of multiple LST-1 protein isoforms that result from multiple annotated transcripts of lst-1, which differ at the 5’ end.

**Figure S2:**
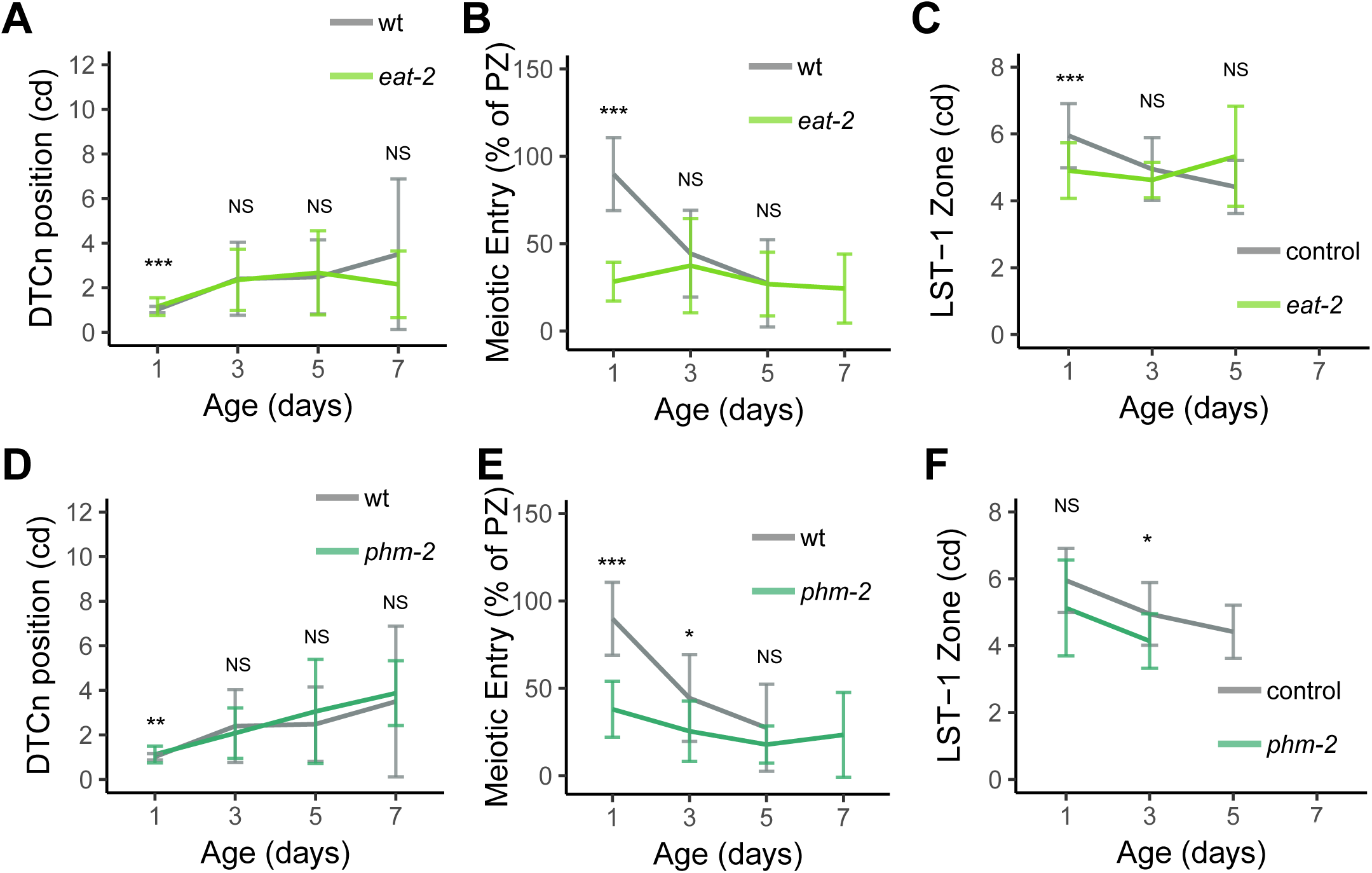
Reproductive and germline phenotypes caused by *eat-2* and *phm-2* mutations. Wild type (gray), *eat-2(ad465)* (light green), and *phm-2(am117)* (dark green) hermaphrodites were mated to males. A,D) Values are the distance of the distal tip cell nucleus (DTCn) from the distal tip measured in cell diameters (cd). B,E) The extent of meiotic entry (measured in cell diameters) expressed as a percent of the extent of the progenitor zone (measured in cell diameters). C,F) The extent of LST-1 in cell diameters (cd) measured from the distal tip to the last row with half or more LST-1 positive cells. The genotypes were: control – lst-1(am302flag); *eat-2* - lst-1(am302flag); eat-2(ad465); *phm-2* - lst-1(am302flag); *phm-2(am117)*. (A-F) Kruskal–Wallis test. * P<0.05, ** P<.001, *** P<.0001

**Figure S3:**
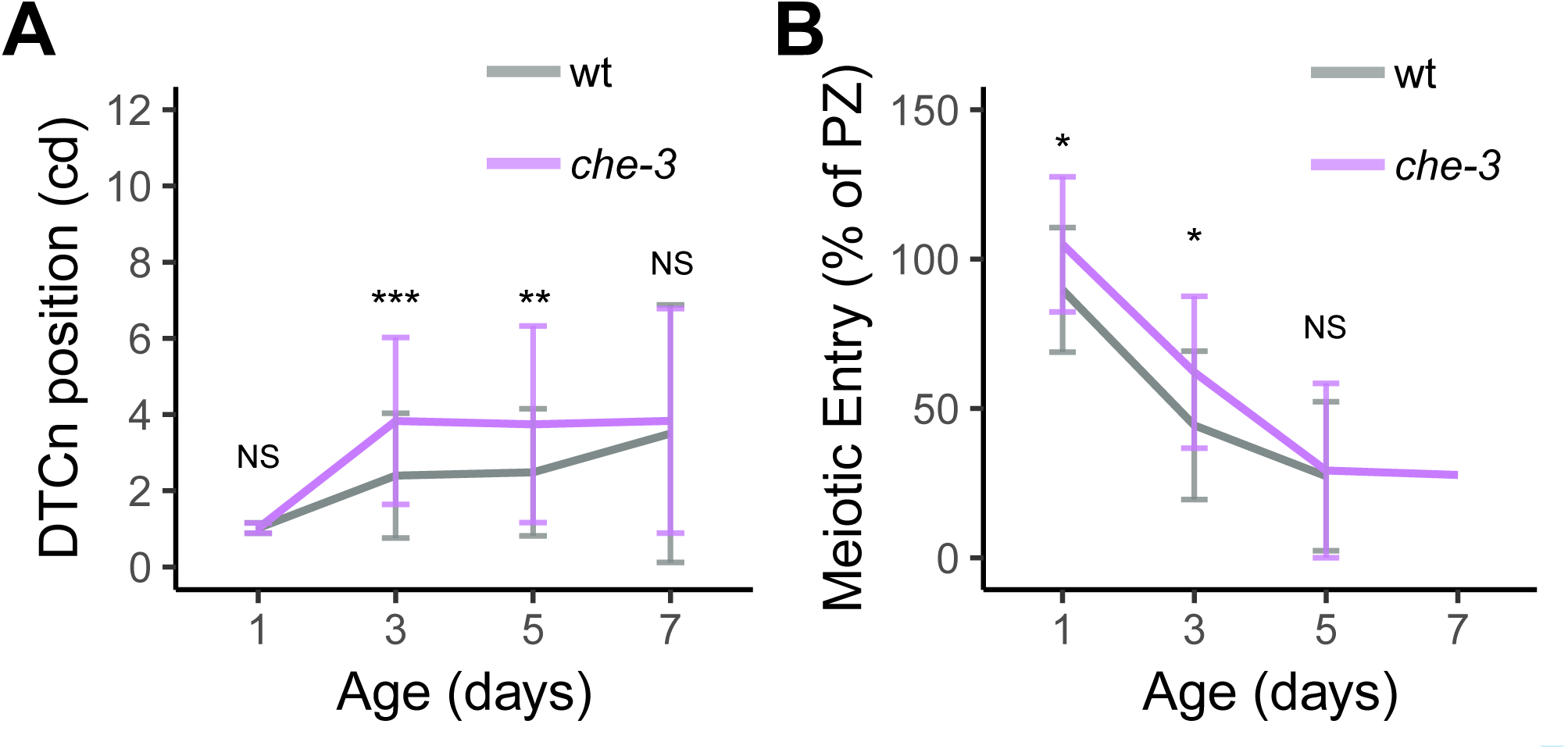
Additional germline phenotypes caused by che-3. Wild type (gray) and *che-3(p801)* (purple) hermaphrodites were mated to males. A) Values are the distance of the distal tip cell nucleus (DTCn) from the distal tip measured in cell diameters (cd). B) The extent of meiotic entry (measured in cell diameters) expressed as a percent of the extent of the progenitor zone (measured in cell diameters). A,B) Kruskal–Wallis test. * P<0.05, ** P<.001, *** P<.0001.

**Figure S4:**
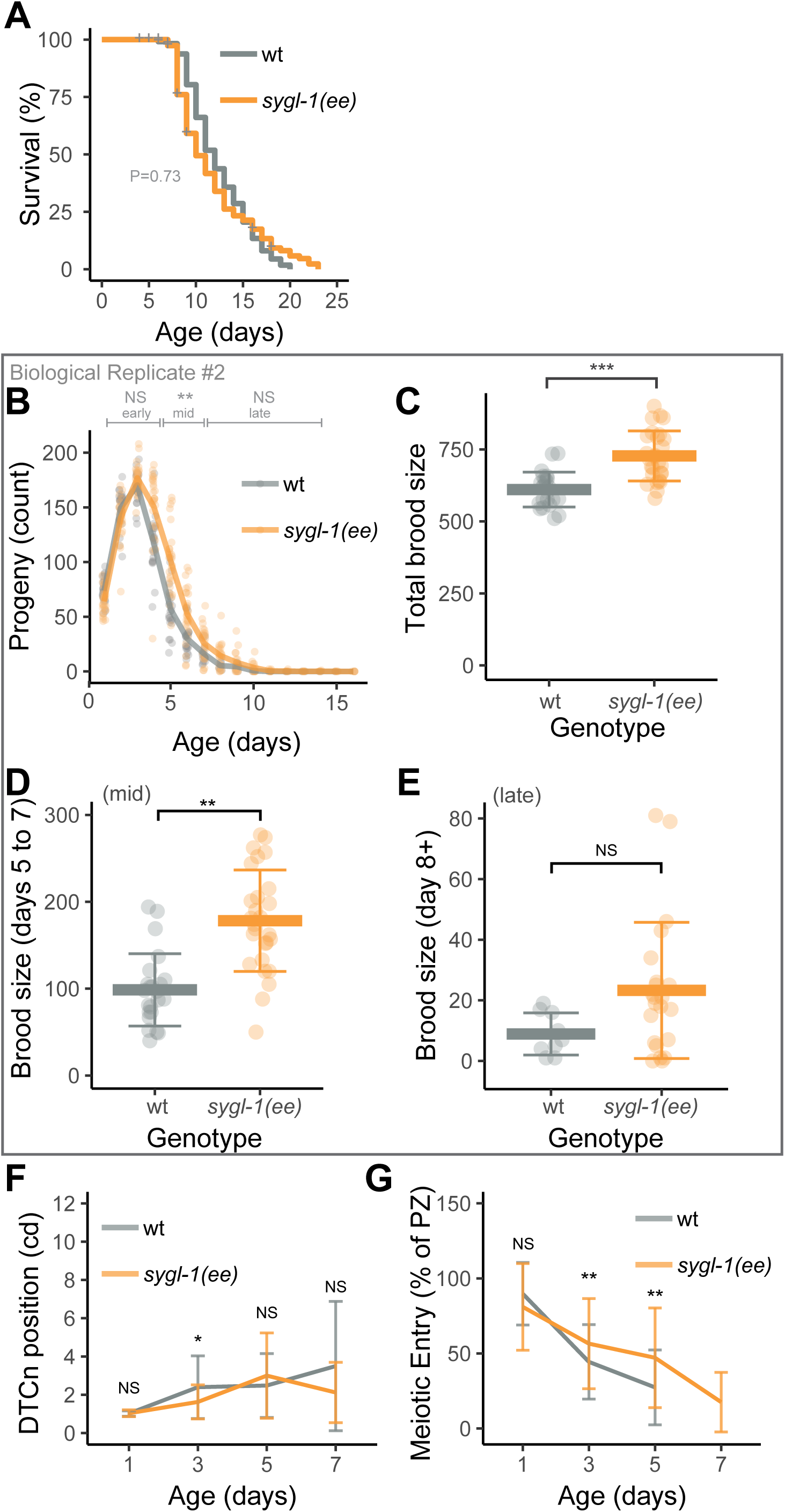
Reproductive and germline phenotypes caused by ectopic expression of SYGL-1. Wild type (gray) and *sygl-1*(ee) (orange) hermaphrodites were mated to males. Genotype of the sygl-1(ee) strain (JK5500) is *sygl-1*(q828null), mut-16(unknown) I; unc-119(ed3lf); qSi150[p*sygl-1*::3xflag::sygl-1::tbb-2 3’utr; Cbr-unc-119(+)] II. A) Kaplan–Meier survival plots analyzed using the Log-rank test. ‘+’ on survival plots indicate animals were censored due to matricidal hatching or crawling off the dish. B) Lines are average daily progeny production, and points are data for individuals. Brackets above indicate early (days 1-4), mid-life (days 5-7) and late (days 8-16) progeny production periods compared by a Wilcoxon Rank Sum test. C) The total brood size, sum of progeny on days 1-15, compared by a Wilcoxon Rank Sum test. D,E) The mid-life and late brood size. F) Values are the distance of the distal tip cell nucleus (DTCn) from the distal tip measured in cell diameters (cd). G) The extent of meiotic entry (measured in cell diameters) expressed as a percent of the extent of the progenitor zone (measured in cell diameters). C-G) Kruskal–Wallis test. * P<0.05, ** P<.001, *** P<.0001.

### Supplemental Table 1: Daily progeny production in the genus Caenorhabditis

#### Footnotes

1. Strain designations – N2 is the standard *C. elegans* wild type. Strains are wild type except fog-2(oz40). Hermaphrodites were self-fertile or mated to conspecific males, and females were mated to conspecific males.
2. Number of mothers analyzed, and total number of progeny.
3. Average number of progeny per mother and standard deviation, after censoring animals due to matricidal hatching.
4. Average number of progeny per mother on adult days 1-16. NA indicates not applicable, as no animals were alive at that age (due to matricidal hatching).

### Supplemental Table 2: Early, mid-life, and late-life progeny production in *C. elegans* mutants

#### Footnotes

1. Genotype. Strain designations are followed by wild type (wt) or gene and allele. Note that the complete genotype of JK5500 is *sygl-1*(q828null), mut-16(unknown) I; unc-119(ed3lf); qSi150[p*sygl-1*::3xflag::*sygl-1*::tbb-2 3’utr; Cbr-unc-119(+)] II.
2. Peak. The number of progeny produced on the day with the largest number. Values are the mean.
3. Early, mid-life, and late-life brood size. The sum of progeny produced on days 1-4 (early), 5-7 (mid-life) and 8-cessation of progeny production (late). Values are mean and standard deviation. P-value from Wilcoxon Rank Sum test is compared to the paired wt control. Red shading indicates P<0.05. Green shading indicates the mutant value is higher than the paired wt control. n= number of hermaphrodites, which is typically lower on days 8+ due to senescent death. In some experiments, only the late brood size was measured.
4. Total brood size. The sum of progeny produced on days 1-cessation of progeny production. Values are mean and standard deviation. P-value from Wilcoxon Rank Sum test is compared to the paired wt control. Red shading indicates P<0.05. Green shading indicates the mutant value is higher than the paired wt control. n= number of hermaphrodites. In experiments where only the late brood size was measured, the total brood size was not determined.

### Supplemental Table 3: Statistical analysis of germline phenotypes, including meiotic entry

#### Footnotes

1. Age. Adult age in days.
2. Progenitor zone. The length of the progenitor zone in cell diameters (cd), measured from the distal tip to the last row with half or more WAPL-1 positive nuclei. Values are mean and standard deviation. P-value from Kruskal-Wallis rank sum test with Dunn post-hoc (non-parametric). All comparisons are to WT of the same age, except day 7 mutant values are compared to day 5 WT values, as sample size was insufficient in day 7 WT. Green shading indicates P<0.05
3. Meiotic entry. The extent of meiotic entry in cell diameters (cd), measured from the end of the progenitor zone to the last EdU+ nucleus following a 10 hour EdU label.
4. Relative meiotic entry. The extent of meiotic entry (measured in cell diameters) expressed as a percent of the extent of the progenitor zone (measured in cell diameters).
5. 5, DTC nucleus position. Values are the distance of the distal tip cell nucleus (DTCn) from the distal tip measured in cell diameters (cd).
6. SYGL-1 expression. The extent of SYGL-1 in cell diameters (cd) measured from the distal tip to the last row with half or more SYGL-1 positive cells.
7. Genotype: control indicates genotype *sygl-1*(am307flag). All other genotypes include sygl-1*(am307flag)* in addition to the listed allele(s).
8. LST-1 expression. The extent of LST-1 in cell diameters (cd) measured from the distal tip to the last row with half or more LST-1 positive cells.
9. Genotype: control indicates genotype lst-1(am302flag). All other genotypes include lst-1(am302flag) in addition to the listed allele(s).

## Materials and Methods

### Strains and General Methods

*C. elegans* strains were cultured at 20°C on 6 cm Petri dishes containing nematode growth media (NGM) agar and a lawn of E. coli strain OP50 unless otherwise noted. The wild-type *C. elegans* strain and parent of edited strains was Bristol N2 (Brenner, 1974). Males of the strain CB4855 (“Mr. Vigorous”) were used to generate mated hermaphrodites because they display a higher mating ability; they also deposit a copulatory plug after mating (Hodgkin and Doniach, 1997). Strains were obtained from the CGC unless otherwise noted. Animals were synchronized by picking fourth-stage larvae (L4) using a dissecting microscope, defined as day 0, from populations that had not experienced starvation for a minimum of three generations.

To obtain populations of mated middle-aged hermaphrodites, we cultured 30-50 L4 hermaphrodites on a dish with 30-50 young adult males (at a 1:1 ratio) for 24 hours, then the hermaphrodites were removed from the males. The presence of copulatory plugs on many of the hermaphrodites confirmed a high frequency of mating in the population; however, hermaphrodites without a copulatory plug were not excluded from the experiment. Prior experiments showed this mating protocol provides sufficient sperm without causing unnecessary damage from male exposure (Kocsisova et al., 2019; Pickett et al., 2013). Hermaphrodites were moved to fresh NGM+OP50 dishes daily until they reached the desired age.

### Progeny Production

To measure progeny production of mated hermaphrodites, we placed 1 L4 hermaphrodite of the appropriate genotype and 3 young adult, CB4855 males on an individual Petri dish with abundant food. After 24 hours, we removed the males and placed each mated hermaphrodite on a fresh dish. We transferred the parent animal to a fresh dish daily. We incubated the previous dish at room temperature (19-23 °C) for two days to allow progeny to develop to adulthood, then scored the presence/absence of male progeny and the number of progeny produced. Progeny that desiccated on the edge of the dish were included in the analysis. For convenience, progeny dishes were sometimes refrigerated at 4 °C to arrest reproduction prior to counting. This method provides accurate and precise measurements of daily progeny production, consistent with previous findings (Hughes et al., 2007).

To measure late progeny production, parent generation animals were mated in batches of 30-50 and transferred to fresh NGM+OP50 dishes in batches, as described in the general methods section above. After 7 days, individual hermaphrodites were moved to fresh NGM+OP50 dishes daily, and progeny were counted after 2 days. In some experiments, 50 μL of 10 mg/mL palmitic acid dissolved in ethanol was applied to the rim of the dish to reduce fleeing (Fletcher and Kim, 2017).

In experiments with *C. elegans, C. remanei*, and *C. brenneri*, conspecific males were used. We noticed that in *C. brenneri*, when three males were placed with one female on each Petri dish, all males except one nearly always fled the dish within 24 hours. *C. elegans* N2 embryonic lethality was 1.2% (n=171). By contrast, several strains of female/male species exhibited high levels of sterility and embryonic lethality. *C. remanei* SB146 was initially relatively healthy, with a brood size average of 390 ± 270, but deteriorated within a few generations from receipt from the *Caenorhabditis* Genetics Center. SB146 displayed ∼75% (n=319) embryonic lethality. *C. brenneri* CB5161 was very sickly with a brood size average of 15 ± 33, and half of the animals displayed full sterility. The average lifespan was 14.5 ± 6 days, and embryonic lethality was higher than 90%. This is probably due to balanced lethal or sterile recessive polymorphisms, identified in genome sequencing (Eric Haag, personal communication). *C. brenneri* VX02253 displayed ∼55% (n=807) embryonic lethality. Even accounting for animals with delayed development, lethality was ∼20% (n=515) when animals were scored after 4 hours, and many progeny that hatched late remained arrested in the L1 stage. Broods from individual hermaphrodites showed a range of lethality ranging from 8% to 55%. Whereas hermaphrodite/male species such as *C. elegans* are well-suited for laboratory culture of small populations, female/male species such as *C. remanei* and *C. brenneri* experience a severe inbreeding depression that poses a challenge to laboratory studies of aging.

Hermaphrodites occasionally failed to mate in these studies, making it possible to measure self-fertile reproduction. The self-fertile brood size of *sygl-1*(ee) hermaphrodites was significantly higher in both biological replicates (262±41 and 306±23 self-progeny in wild type; 382±47 and 476±91 self-progeny in *sygl-1*(ee); P=0.0025 and 0.013, respectively). Because self-fertile brood size is usually determined by the number of sperm produced during development, this result suggests that *sygl-1*(ee) hermaphrodites produce more self-sperm than wild type, a phenotype known as partial <u>m</u>asculinization <u>o</u>f the <u>g</u>ermline (Mog).

### Lifespan measurements

Studies of lifespan were begun on day zero by placing ∼60 L4 stage hermaphrodites on a Petri dish. Hermaphrodites were transferred to a fresh Petri dish daily during the reproductive period (approximately the first ten days) to eliminate self-progeny and every 2-3 days thereafter. Each hermaphrodite was examined daily using a dissecting microscope for survival, determined by spontaneous movement or movement in response to prodding with a platinum wire. Dead worms that displayed matricidal hatching, vulval extrusion, or desiccation due to crawling off the agar were excluded from the data analysis. Data is aggregated from 2-3 simultaneous biological replicates starting with up to 60 worms each. Mean lifespan and the log-rank test were calculated using OASIS 2 (Han et al., 2016).

### Embryonic viability

Individual young adult hermaphrodites were placed on a dish and allowed to deposit eggs. After several hours, the eggs were picked to a fresh dish and arranged in a line next to the lawn of E. coli OP50. After 16 hours, the line was observed and any unhatched eggs were counted. After 48 hours, the line was observed again, and eggs which still had not hatched were counted.

### EdU labeling experiments

To make EdU dishes, we seeded M9 agar dishes (Stiernagle, 2006) with concentrated E. coli MG1693 Thy-which had been grown for 24 hours at 37C with shaking in minimal media containing 20uM 5-ethynyl-2’-deoxyuridine (EdU, Invitrogen). The culture consisted of 100 mL M9, 4 ml overnight LB-grown MG1693 E. coli, 5 ml 20% glucose, 50 μl 1.25 mg/ml thiamine, 1.2 ml 0.5 mM thymidine, 100 μl 1M MgSO _4_, 200 μl 10mM EdU (Fox et al., 2011).

Appropriately mated and aged animals were either washed with phosphate buffered saline (PBS) or picked to EdU dishes and incubated for the appropriate time: 0.5, 4, 7, or 10 hours, then dissected and processed as detailed in the following section. Nuclei with any amount of EdU signal overlapping with DAPI staining were scored as EdU-positive. For the full detailed protocol, see Kocsisova et al. (2018).

### Dissection and Immunohistochemistry

The dissection and staining protocol follows the batch method (Francis et al., 1995), where all dissected tissues were incubated within small glass tubes rather than on a slide. Animals were washed with phosphate-buffered saline (PBS) into a dissecting watchglass (Carolina Biological Item # 742300), immobilized with Levamisole (final concentration 200uM), and dissected with a pair of 25G 5/8” needles (PrecisionGlide from BD) by cutting at the pharynx and/or at the tail. Dissected gonads were fixed in 2 mL 3% paraformaldehyde (PFA) (10 mL 16% PFA, EM Grade, Electron Microscopy Sciences, Hatfield, PA Catalog No 15710) phosphate-buffered solution for 10 minutes at room temperature and then post-fixed with 2 mL 100% methanol (Gold-label from Fisher) at - 20C for 1 hour or longer (up to several days).

Fixed gonads were rehydrated and washed three times in PBS + 0.1% Tween-20 (PBSTw), then incubated in 100 μl of primary antibody at room temperature for 4-24 hours, washed 3 times in PBSTw, and incubated in 100 μl of secondary antibody at room temperature or 4C for 2-24 hours. Antibodies were diluted in 30% goat serum (Gibco C16210-072) in PBS. Primary antibodies used were: rabbit-anti-WAPL-1 (Novus Biologicals Cat#49300002, Lot G3048-179A02) (1:2000), mouse-anti-MSP (Major Sperm Protein) (Miller et al., 2001) (1:2000), mouse-anti-pH3 (Millipore clone 3H10 Cat#05-806, Lot#2680533) (1:500), mouse-anti-FLAG (SIGMA M2) (1:1000). Secondary antibodies were: goat-anti-mouse IgG-conjugated Alexa Fluor 488/594/647, goat-anti-rabbit IgG-conjugated Alexa Fluor 488/594/647 (Invitrogen).

Following antibody staining, gonads were washed 3 times in PBSTw. An EdU Click-iT reaction was performed according to manufacturer’s instructions (Invitrogen C10350), using 100 μl reagent for a 30 minute incubation, followed by a quick wash in manufacturer-supplied rinse buffer and four ∼15 minute washes in PBSTw. Stained gonads were resuspended in 1 drop of Vectashield containing 4’,6-Diamidino-2-Phenylindole Dihydrochloride (DAPI) (Vector Laboratories H-1200), applied to a large pre-made agarose pad on a glass slide, and covered with a 22 mm x 40 mm #1 cover glass. The slide was allowed to settle overnight at room temperature, sealed with clear nail polish, and stored at 4 °C as needed. Images were acquired within 72 hours when possible.

### Confocal Imaging

Images were collected using a Zeiss Plan Apo 63X 1.4 oil-immersion objective lens on a PerkinElmer Ultraview Vox spinning disc confocal system on a Zeiss Observer Z1 microscope using Volocity software. Approximately twenty 1 μm z-slice images were acquired for each gonad. Images were exported as hyperstack .tif files for further analysis.

### Image analysis

Images were stitched either in Volocity or using the Image J plugins for pairwise stitching and Grid/Collection of sequential images (Preibisch et al., 2009). Images were rotated, cropped, arranged, and annotated in Illustrator (Adobe).

Nuclei were manually counted in each z-slice where they occurred. The person performing counts was not blinded to experimental groups, because the differences were generally obvious to an experienced observer. Counts of nuclei were performed in Fiji/ImageJ (Schindelin et al., 2012) using the Cell Counter plug-in (Rasband, 2016; De Vos, 2015). To remove multiply-counted nuclei, a modified version of the R-script Marks-to-Cells was used (Seidel and Kimble, 2015). This script improved the precision of cell counts in the germline.

#### Nuclear morphology

DAPI staining was used to assess meiotic prophase stages, nuclear counts, row counts and progression of gametogenesis. Endomitotic oocytes in the proximal gonad arm, distal to the spermatheca, were recognized as large DAPI stained blobs. Endomitotic oocytes are known to result from failure to coordinate meiotic maturation of diakinesis stage oocytes with ovulation, resulting in the unfertilized mature oocytes that are mitotic cell cycling without cytokinesis because of the absence of the sperm derived centriole (Greenstein, 2005; Iwasaki et al., 1996; McCarter et al., 1997). This is distinct from endomitotic oocytes in the uterus, which naturally occur in older unmated hermaphrodites that have exhausted their self-sperm. In this case the most proximal oocyte undergoes low frequency spontaneous maturation and ovulation, but because there is no sperm in the spermatheca the matured but unfertilized oocyte begins endomitotic cycling (McCarter et al., 1999). Endomitotic oocytes were identified from dissected germlines; as the proximal germline was not always visible, this raises the possibility that the frequency was under-estimated.

#### Progenitor zone

The PZ has been previously called the mitotic zone or the proliferative zone. We employed staining with the cohesin chaperone WAPL-1 to measure the length of the PZ (Kocsisova et al., 2019; Mohammad et al., 2018). Other studies of unmated day 1 hermaphrodites used the crescent-shaped DAPI morphology of leptotene nuclei to approximate the proximal boundary of the PZ (Crittenden et al., 2006; Roy et al., 2016).

#### SYGL-1 and LST-1

The length in cd and the number of cells in the PZ that contain cytoplasmic SYGL-1 and LST-1 was assessed with anti-FLAG antibody staining using strains where CRISPR genome editing was employed to insert a 3xFLAG epitope onto the endogenous gene product (Kocsisova et al., 2019; Shin et al., 2017).

#### Distal Tip Cell nucleus

The Distal Tip Cell nucleus (DTCn) can be visualized by WAPL-1 staining on the periphery of the gonad. The distal tip cell secretes the LAG-2 Notch ligand, and the distal tip cell nucleus displays an age-related increase in distance from the distal end of the gonad, which reflects mispositioning (Kocsisova et al., 2019). There was no significant difference between wild type and *daf-2(e1370)* in the position of the distal tip cell nucleus at days 1, 3, 5 and 7 (**Figure S1G, Table S3**). The distal tip cell nucleus was shifted further in *che-3(p801)* mated hermaphrodites at days 3 and 5 than in wild-type, but not significantly different at days 1 and 7 (**Figure S3A, Table S3**). The distal tip cell nucleus was shifted to a similar extent in *sygl-1*(ee) mated hermaphrodites compared to wild-type (**Figure S4F, Table S3**).

### Statistical Analysis

Statistical analyses were performed in R (R Core Team, 2013) using R studio (RStudio Team, 2015) and the following packages: ggplot2 (Wickham, 2009), svglite, plyr, and tidyR. Individual measurements and script used to analyze them are provided in supplemental tables and files. Depending on the type of variable, the following tests were performed. For continuous measurement variables, the Kruskal-Wallis Rank-sum test was used with a Dunn post-hoc and p-values adjusted with the Benjamini-Hochberg FDR method. Data were also compared using an ANOVA with a Tukey post-hoc test. Brood sizes were compared using a pairwise Wilcoxon Rank Sum test (also known as a Mann-Whitney U test). For categorical measurement variables, the Pearson’s Chi-squared test of independence was used with post-hoc p-values adjusted with the False Discovery Rate method. Statistical comparisons were made between mutant and wild-type animals of the same age, with the exception noted in the supplemental tables and figure legends. In the text all data are represented as mean ± SD. All error bars shown in figures represent the mean +/- standard deviation. NS indicates P>0.05, * P<0.05, ** P<.001, *** P<.0001.

All data were subjected to consistent exclusion criteria. Animals that displayed evidence of matricidal hatching or sperm depletion (as indicated by lack of male progeny or lack of Major Sperm Protein immunofluorescence) were excluded from all analyses unless otherwise noted. Animals that displayed sporadic phenotypes such as endomitotic oocytes, a shifted distal tip cell, or failure to exhibit any EdU staining were excluded from analyses. Animals which fled the plate are excluded from the brood size calculations. Animals that died by matricidal hatching are included in the brood size calculations. For this reason, it is possible that brood sizes are slight under-estimates. Matricidal hatching was very common, and excluding these animals would have biased the dataset.

## Acknowledgements

We are grateful to the E. coli stock center for MG1693; Wormbase; the *Caenorhabditis* Genetics Center which is funded by the National Institutes of Health Office of Research Infrastructure Programs (P40OD010440) for strains; Charles Baer for VX02253; the Kimble lab for JK5500; Aiping Feng for reagents; Andrea Scharf, Sandeep Kumar, Ariz Mohammad, and John Brenner for training, advice, support, reagents, and helpful discussion; and Andrea Scharf and Aaron Anderson for feedback on this manuscript.

## Authors’ contributions

ZK, KK, and TS designed the study.

ZK, EB, JS, HCL, and DS performed the experiments.

ZK, EB, JS and HCL analyzed data.

ZK wrote the manuscript.

ZK, KK, and TS revised the manuscript.

All authors read and approved the final manuscript.

## Abbreviations Used

(PBS): Phosphate Buffered Saline
(DAPI): 4’,6-Diamidino-2-Phenylindole, Dihydrochloride
(EdU): 5-ethynyl-2’-deoxyuridine
(DTC): Distal Tip Cell
(DTCn): Distal Tip Cell nucleus
(PZ): Progenitor Zone
(UTR): untranslated region
(cd): cell diameters

